# Epistasis, aneuploidy, and gain-of-function mutations underlie the evolution of resistance to induced microtubule depolymerization

**DOI:** 10.1101/2020.10.07.327759

**Authors:** Mattia Pavani, Paolo Bonaiuti, Elena Chiroli, Fridolin Gross, Federica Natali, Francesca Macaluso, Adam Poti, Sebastiano Pasqualato, Simone Pompei, Marco Cosentino-Lagomarsino, Giulia Rancati, David Szuts, Andrea Ciliberto

**Affiliations:** IFOM, The Firc Institute of Molecular Oncology, Via Adamello 16, 20139, Milano, Italy; Institute of Medical Biology (IMB), Agency for Science, Technology and Research (A*STAR), Singapore 138648, Singapore; IEO, European Institute of Oncology IRCCS, Via Adamello 16, 20139 Milan, Italy; Institute of Enzymology, Research Centre for Natural Sciences, H-1117 Budapest, Hungary

## Abstract

Microtubules, polymers of alpha- and beta-tubulin, are essential cellular components. When microtubule polymerization is hindered, cells are delayed in mitosis, but eventually they manage to proliferate with massive chromosome missegregation. Several studies have analyzed the first cell division upon microtubules impairing conditions. Here, we asked how cells cope on the long term. Taking advantage of mutations in beta-tubulin, we evolved in the lab for ∼150 generations 24 populations of yeast cells unable to properly polymerize microtubules. At the end of the evolution experiment, cells re-gained the ability to form microtubules, and were less sensitive to microtubule depolymerizing drugs. Whole genome sequencing allowed us to identify genes recurrently mutated (tubulins and kinesins) as well as the pervasive duplication of chromosome VIII. We confirmed that mutations found in these genes and disomy of chromosome VIII allow cells to compensate for the original mutation in beta-tubulin. The mutations we identified were mostly gain-of-function, likely re-allowing the proper use of the mutated form of beta-tubulin. When we analyzed the temporal order of mutations leading to resistance in independent populations, we observed multiple times the same series of events: disomy of chromosome VIII followed by one additional adaptive mutation in either tubulins or kinesins. Analyzing the epistatic interactions among different mutations, we observed that some mutations benefited from the disomy of chromosome VIII and others did not. Given that tubulins are highly conserved among eukaryotes, our results are potentially relevant for understanding the emergence of resistance to drugs targeting microtubules, widely used for cancer treatment.

## INTRODUCTION

Microtubules are essential components of the cytoskeleton, formed by alpha- and beta-tubulins (Tub1/3 and Tub2 in budding yeast) [1, 2]. During mitosis, they form the mitotic spindle which is responsible for the segregation of sister chromatids to the daughter cells. Microtubules of the mitotic spindle via a process of ‘search and capture’ interact with chromosomes at specialized proteinaceous structures called kinetochores [3]. As long as there are unattached or improperly attached kinetochores, cells are arrested in pro-metaphase by a signaling pathway called the mitotic checkpoint or spindle assembly checkpoint (SAC) [4]. After all chromosomes are properly attached, the checkpoint is silenced, and cells transit into anaphase.

Microtubule dynamics plays a key role both for the search and capture of chromosomes, and for segregating them to the opposite poles of the cell. As cells need to carefully modulate microtubule dynamics, the latter depends on several different factors. Tubulins can polymerize or shrink, the alternation between the two being heavily affected by the status of the GTP bound to beta-tubulin on the plus-end of the filament. At centrosomes, gamma-tubulins contribute to polymerization by nucleating new filaments. In budding yeast, this requires the gamma-tubulin small complex (gamma-TuSC), which is formed by gamma-tubulin, Tub4 in budding yeast, and two co-factors called Spc98 and Spc97 [5]. Tub4 also requires GTP for microtubules polymerization [6]. Besides tubulins themselves, other proteins interact with microtubules and control their polymerization. Among them are kinesins, specialized motors that move along filaments, some of which can also depolymerize microtubules [7]. In budding yeast, Kip3, which belongs to the kinesin-8 family, is such a kinesin with depolymerization activity [8, 9].

Tubulins are coded by essential genes, highly conserved among eukaryotes. Even mutations that partially impair their function have quite dramatic effects. Early screens in yeast identified a wide array of temperature-sensitive mutations that either depolymerize or hyperstabilize microtubules [10]. Cells expressing these alleles are very sick and delayed in mitosis. Similarly, drugs that interfere with microtubule polymerization (stabilizers -- eg taxanes -- or destabilizers -- eg vinca alkaloids) affect chromosome-microtubule attachment, activate the mitotic checkpoint and arrest cells before anaphase. By delaying cell cycle progression, microtubule drugs can promote apoptosis in transformed cells [11].

In the long-term, the effects of antimitotics are jeopardized by the emergence of resistance. Mechanisms of resistance have been studied, including the differential expression of different isotypes of tubulin, expression levels of microtubule-associated proteins, and multi drug resistance mechanisms [12] [13] [14]. The role of mutations in tubulin genes has been debated. They develop frequently in cell lines, but have been harder to find in patients [15]. Recent data, however, have yet again pointed at a potential role for gain-of-function mutations in tubulin genes [16]. Interestingly, mutations found in patients and tested in cell lines were shown to have an effect on microtubule dynamics. In this context, it was underlined the potential role for tubulin mutations not generally in cancer development but rather as an adaptive response to drugs [17]. Finally, given the multi-layered control of microtubule stability, not surprisingly mutations in proteins interacting with tubulin such as kinesins have also been reported to affect the development of resistance to drug treatment [18, 19].

Although mutations related to resistance against microtubule drugs have been identified, basic questions still have no answer. Is there a typical sequence of mutations that gives rise to resistance (ie, is the process of resistance repeatable and thus potentially predictable [20])? Adaptive mutations that restore essential functions are often loss-of-function [21, 22]. Is this also the case for tubulin? Do aneuploidies play a role in resistance? The question is relevant since drugs affecting microtubule-kinetochore attachments are likely to cause chromosome missegregation, and aneuploidies have been shown multiple times to be adaptive in stressful conditions [22-25]. However, in mammals a role for aneuploidy in the development of resistance to induced microtubules depolymerization has not been reported yet.

These questions require approaching the emergence of resistance from an evolutionary viewpoint [23, 25-28]. This approach, which requires genetically well-characterized systems, allows deciphering the dynamics of genetic changes underlying the emergence of resistance. Importantly, lab evolution experiments also can shed light on the epistatic interactions among adaptive mutations, and thus on possible mechanisms for avoiding or delaying the emergence of resistance [28]. Here, we performed a laboratory evolution experiment to study how cells develop resistance to forced microtubule depolymerization. We used haploid yeast cells expressing an allele of beta tubulin (*tub2-401*) which carries four point mutations that result in three amino acid changes. This is a cold-sensitive allele, which cannot polymerize microtubules when grown at low temperature [29, 30]. Several other alleles are available, but *tub2-401* is the one showing the most penetrant phenotype. We opted for a mutation mimicking the effect of drugs, rather than drugs themselves (eg, benomyl, nocodazole), to avoid the development of resistance via generic multi-drug mechanisms [31].

We grew cells for more than one hundred generations at the semi-restrictive temperature, until they recovered growth. We confirmed that evolved cells re-gained the ability to assemble regular spindles, and we identified two pathways through which cells can acquire resistance. In both of them disomy of chromosome VIII (*chrVIII 2X*) is the likely initial step. Our results may be relevant for understanding the emergence of resistance in higher eukaryotes as well.

## RESULTS

### Yeast cells become resistant to stimuli inducing microtubules depolymerization

We evolved haploid yeasts carrying the *tub2-401* allele (Figure S1A). At low temperature, these cells are unable to properly polymerize microtubules [29, 30] and activate the mitotic checkpoint [32]. We grew them at an intermediate temperature (18 °C), where cells can still proliferate, albeit less efficiently than wild types. For each genotype, we evolved multiple populations (8 wild types and 24 *tub2-401*), each started from an individual clone of the same ancestor (Figure 1A). Measurements of growth started after one day at 18 °C (Generation 0 or G0 -- Figure 1B).

**Figure 1.**
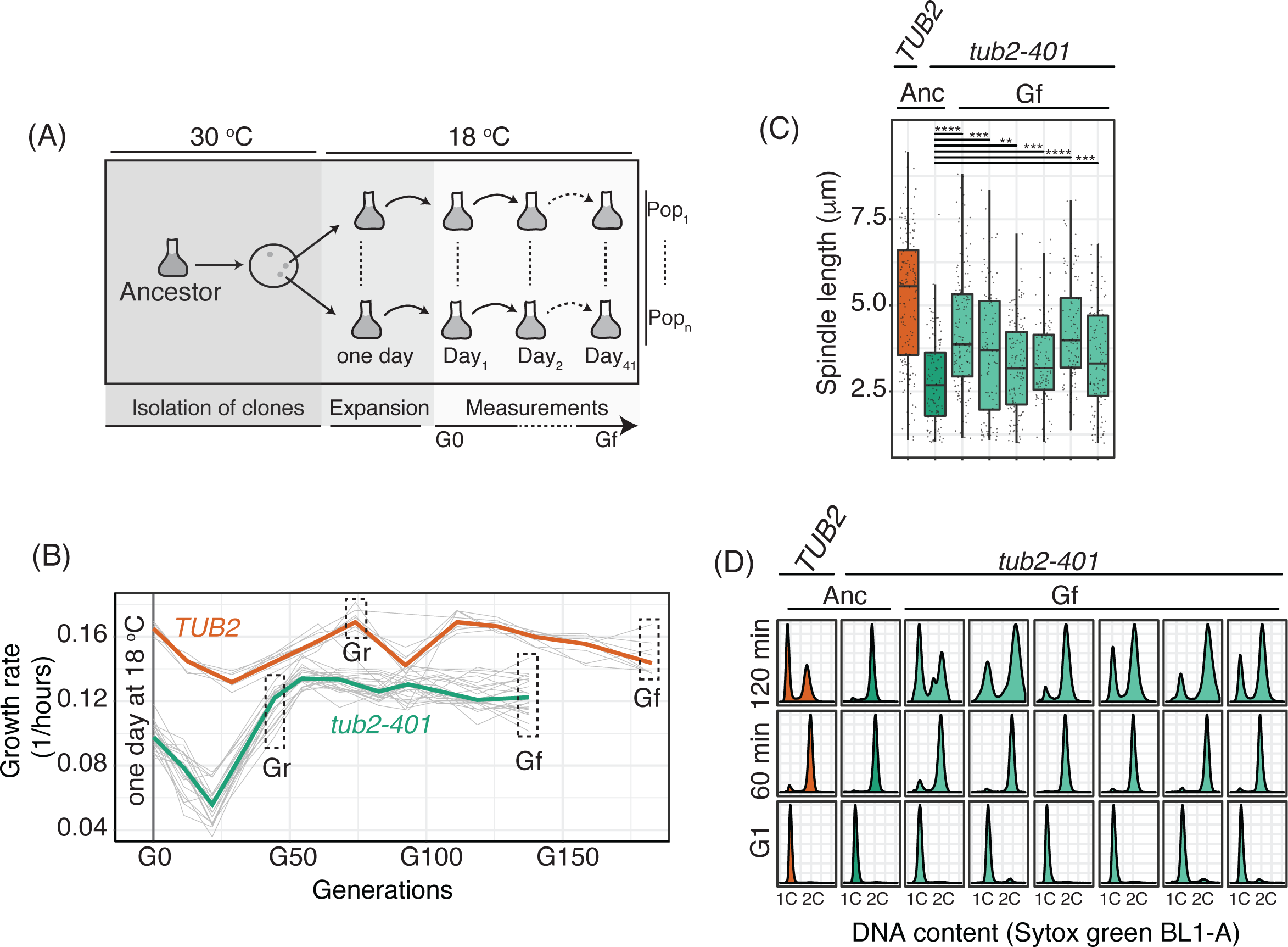
Cells impaired for microtubule polymerization increase their growth rate during a laboratory evolution experiment. a) Scheme of the evolution experiment. From the clonal ancestor, we produced 8 populations of *TUB2* cells, and 24 populations of *tub2-401*. The first measurement of growth rate was done after one day at 18 °C (generation 0 or G0). The last measurement was done at generation final (Gf). Between the two, we also analyzed by NGS cells immediately after recovery of growth (generation recovery or Gr). b) Every 3-4 days the growth rates of the different populations were measured. Here, growth rate is the net output between cell division and cell death. c) Ancestral cells and selected populations of cells collected at the end of the experiment (Gf) were synchronized in G1 at 30 °C and released at 18 °C. Tub1 was detected by immunofluorescence while nuclei were stained with DAPI. Spindle lengths were measured with a custom software across 3 timepoints centered on the maximum proportion of large-budded cells (dumbell) (Figure S1C). For selecting evolved populations, we chose those carrying only one mutation with high frequency in the three most frequently mutated genes -- *KIP3, TUB2*, or *TUB1* (populations G5, D4, B4, D5, H6, E5) (Figure 3A). d) Ancestral cells and selected Gf populations (G5, D4, B4, D5, H6, E5) were thawed at 30 °C and cultured overnight. Cells were synchronized in G1 and released in nocodazole (2 μg/ml, re-added after 2.5 hours at 1 μg/ml). Cells were collected after 1 and 2 hours after nocodazole addition. DNA content was assessed by Sytox Green staining.

Cells expressing *tub2-401* grew slower than control cells expressing wild type *TUB2* (Figure 1B). After ∼20 generations, the growth rate (a combined effect of cell division and cell death) started to increase, and after ∼45 generations approached the wild type. It fluctuated around the same value for another ∼100 generations, when we stopped the experiment (final Generation or Gf -- Figure 1B).

At low temperature, cell cycle progression of *tub2-401* is delayed in mitosis [32]. Regardless of the prolonged mitotic arrest, many cells slip through and continue proliferating. Cell division, however, takes place when microtubules are not properly polymerized and thus chromosomes tend to missegregate [32], which we confirmed by following the inheritance of GFP-tagged chromosome V by live cell imaging during the first cycle after decreasing temperature (Figure S1B). Hence, we interpreted the initial slow growth rate as a consequence of prolonged activation of the mitotic checkpoint. Likely, cell death caused by massive chromosome missegregation contributes to the apparent initial decrease of growth.

To explain how cells recovered growth, we hypothesized that some of them acquired mutations allowing them to assemble more structured mitotic spindles that segregated chromosomes more efficiently. As such, the mitotic checkpoint was lifted, and fitter cells progressed more rapidly. To test this interpretation, we analyzed both mitotic spindles and checkpoint response in cells carrying the *tub2-401* mutations at Gf (end of the experiment, Figure 1B). To analyze mitotic spindles, a subset of evolved populations was synchronized in G1 by alpha-factor and released at 18 °C. We then measured the length of spindles across three different time points, centered on the time when the fraction of cells with a large bud (a feature of mitotic arrest) was the highest (Figure S1C). Our data show that evolved cells have longer spindles than ancestors, but shorter than wild types (Figure 1C, Figure S1D).

As a confirmation that evolved cells spend less time in mitosis, we analyzed their size (for which we used FSC measured by FACS). During the first cycles under checkpoint-activating conditions, *tub2-401* cells increase their size, due to the prolonged arrest in mitosis. Thus, average size can be used as a proxy for checkpoint activation. [32]. Accordingly, we observed that ancestors *tub2-401* were larger than controls expressing *TUB2*. At the end of the experiment, instead, their size approached that of evolved controls (Figure S1E).

The enhanced ability of evolved cells to polymerize microtubules was not uniquely related with overcoming the *tub2-401* mutations, but was confirmed also upon treatment with microtubules depolymerizing drugs. After synchronization in G1 and release in low concentration of nocodazole at 30 °C, evolved cells kept the arrest for a shorter time than their ancestors. At this concentration, wild type cells do not arrest, whereas *tub2-401* cells are delayed in mitosis due to increased sensitivity to microtubule depolymerization. Evolved *tub2-401* showed less sensitivity to nocodazole, in-between ancestors and wild type cells (Figure 1D). Under higher concentration of nocodazole, all strains mounted an efficient checkpoint response (not shown).

In conclusion, we showed that evolved cells became less sensitive to induced microtubule depolymerization, either caused by the *tub2-401* allele or nocodazole.

### Evolved strains mutate recurrently a small set of genes

With the aim of understanding the evolutionary dynamics, we addressed the genetic basis of resistance. We sequenced all populations at the final generation, and looked for genes that were mutated more than once in different populations with unique mutations. We then collected all mutations occurring in these recurrently mutated genes. Hereafter, we focused our attention only on these mutations (Table S1).

Control cells did not experience impairment of microtubule polymerization, yet they were under stress due to the low temperature. In these cells, genes of the *PHO* pathway (*PHO4* and *PHO81*, Figure S2A-B) were recurrently mutated. We did not observe any change of ploidy (Figure S2C). In cells expressing *tub2-401*, we identified recurrently mutated genes that were obviously related with microtubules dynamics, and primarily tubulins themselves. Several mutations affected *TUB2* (Figure 2A). They were all missense mutations, in agreement with *TUB2* being essential in budding yeast. We also observed mutations in *TUB1* (alpha-tubulin) and components of the gamma-tubulin complex (*SPC98*). Like *TUB2*, these essential genes had missense mutations (Figure 2A). Besides tubulins, we found mutated multiple times the gene coding for the kinesin-8 Kip3. Here, mutations were not only missense, but also nonsense and frameshifts, spread all over the gene (Figure 2B). Finally, we observed disomy of chromosome VIII (*chrVIII 2X*) in the large majority of populations carrying *tub2-401* (Figure 2C, Figure S2D). In only one population, we observed disomy of chromosome III. We did not identify recurrent structural variants.

**Figure 2.**
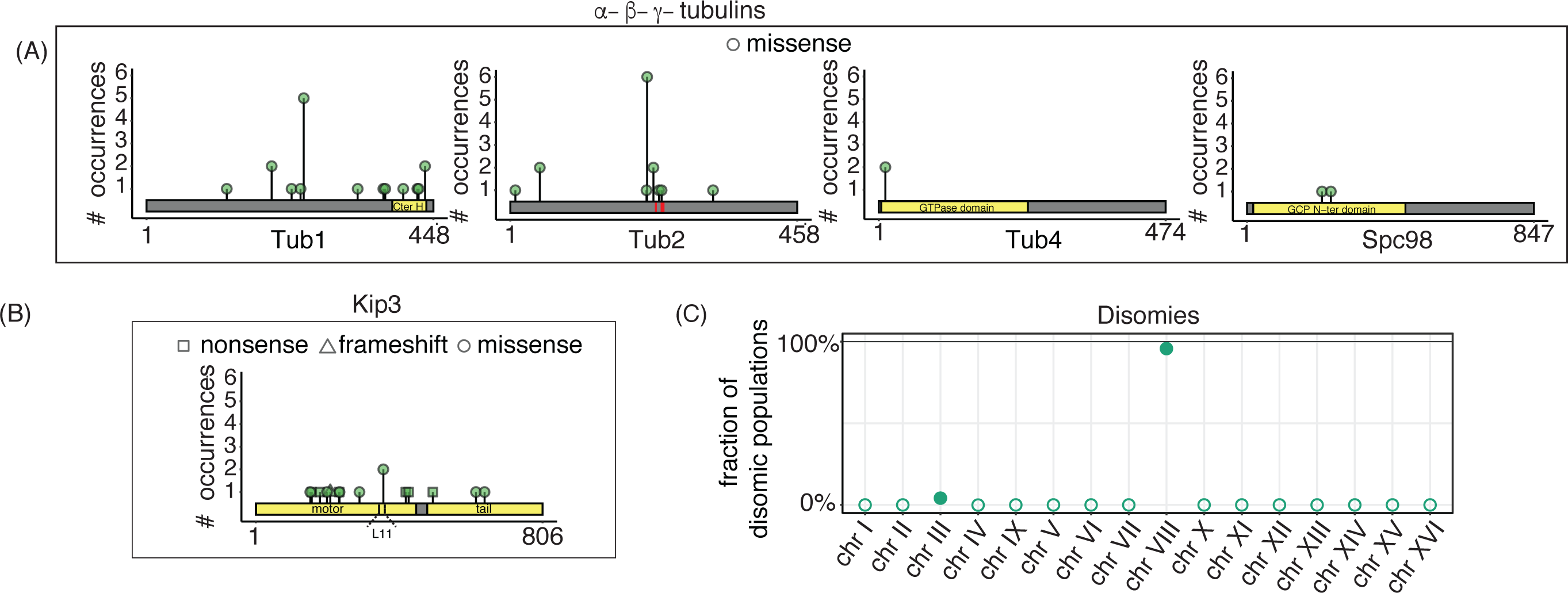
Amino acid changes in recurrently mutated genes. a) Amino acid changes caused by mutations occurring independently multiple times in alpha, beta and gamma tubulin genes. In the analysis, we sum up all mutations that affect the same amino acid residue. Only missense mutations were found. b) Amino acid changes caused by mutations occurring independently multiple times in *KIP3*. c) Fraction of disomic populations. Empty dots are used when there are no disomic chromosomes. Chromosome copy numbers were determined by coverage analysis (an example in Figure S2D). All data were collected at the end of the evolution experiment (Gf in Figure 1B).

Some genes originally identified by our pipeline were not followed up in our analysis. Among them, genes of the *ADE* pathway which have already been reported mutated in evolution experiments of yeast with the same genetic background (W303) but independently from impaired microtubule polymerization and low temperature [33]. We also did not follow up the two genes which have not been characterized yet (*YHR033W, YJL070C*) and *PRR2*, as they are present with very low frequency in two populations only (Table S1). Instead, we followed up *TUB4*, which is mutated twice with high frequency but always with the same mutation, since its protein product interacts directly with Spc98.

In summary, cells unable to properly polymerize microtubules recurrently mutate primarily genes belonging to two classes: i) Tubulins and members of gamma-TuSC (*TUB1, TUB2, TUB4, SPC98*); and ii) *KIP3*. Moreover, we noticed that the large majority of strains carrying the *tub2-401* mutations are disomic for chromosome VIII.

### Cells acquire one mutation in recurrent genes after becoming disomic for chromosome VIII

Next, we analyzed the genetic composition of the individual populations. All of them but one, had at least one recurrently mutated gene (Figure 3A). In several populations we observed the co-presence of multiple mutations, some of them affecting the same gene (Figure S3A). Strikingly, in almost all populations the sum of allele frequencies of mutations approached but did not exceed 100% (Figure 3A). This is not the theoretical maximum, if cells would carry more than one mutation, the sum of frequencies would be higher than 100%. Hence, this result suggests that individual cells are likely to carry only one mutation. To confirm this hypothesis, we extracted and analyzed clones from populations with more than one mutation at high frequency. By Sanger sequencing, we never found two mutations in the same clone (Table S2). The pervasive presence of mutations in the genes we identified among evolved populations, and the fact that the sum of mutation frequencies approaches 100% suggest that the identified mutations are largely responsible for recovered growth.

**Figure 3.**
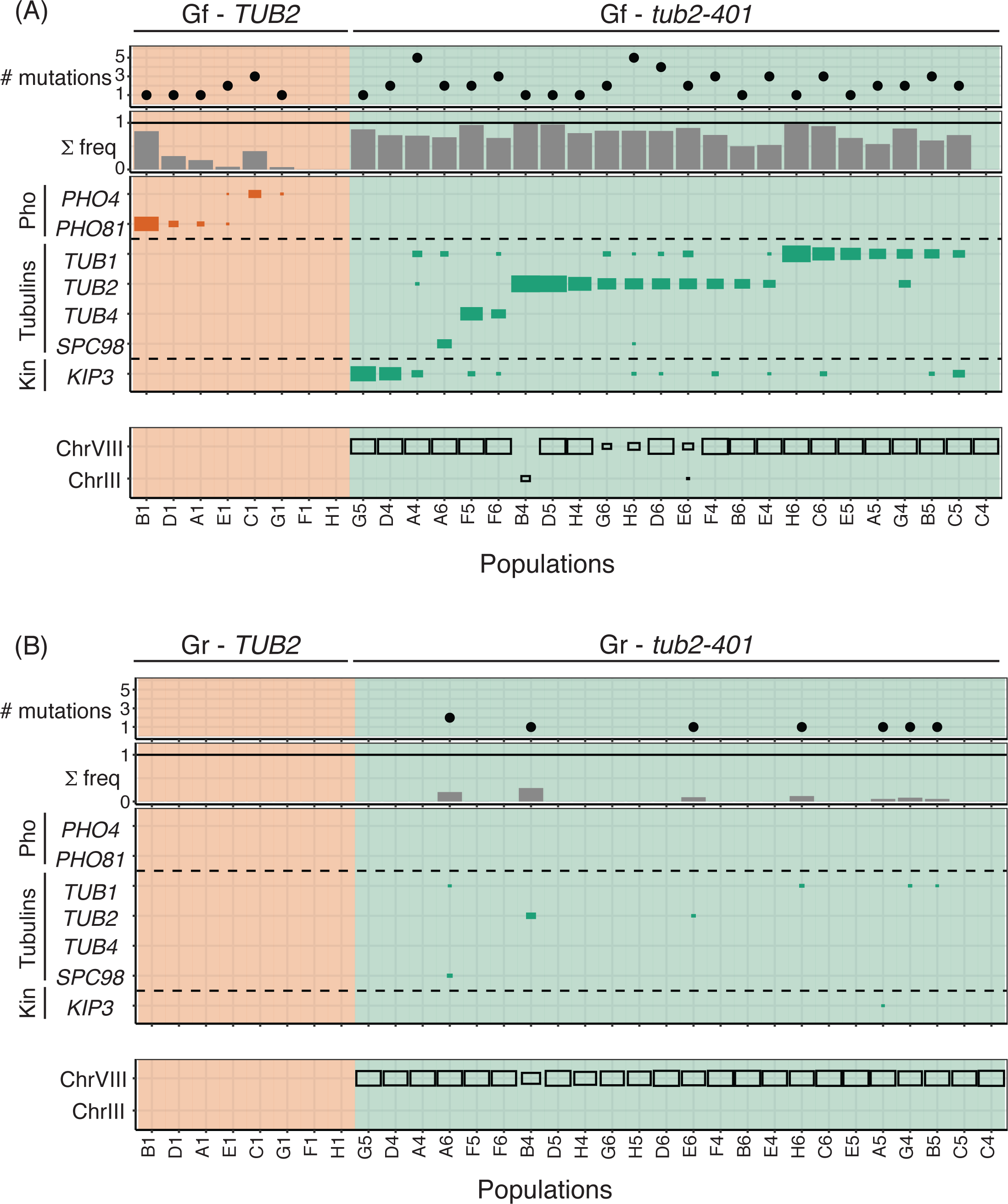
Mutations frequencies measured at intermediate (Gr) and final (Gf) time points. a-b) The tables summarize the number and frequencies of mutations in recurrently mutated genes within each population (column). First panel from top: number of mutations in recurrently mutated genes for each population. Notice that there can be more mutations for the same gene. Second panel from top: sum of the frequencies of all mutations in recurrently mutated genes for each population. Third panel from top: mutation frequency of recurrently mutated genes for each population. In each row we group together frequencies of all mutations affecting one specific gene. The size of squares is proportional to the mutation frequency. Bottom panel: frequency of disomic chromosomes within each population. The size of the squares is proportional to the frequency. In (a), we analyzed cells at the end of the experiment (Gf in Figure 1B), in (b) after cells have started recovering growth (Gr in Figure 1B).

It is striking that regardless of the variability in mutation frequencies, in almost all populations we detected disomy of chromosome VIII only. This result suggests that *chrVIII 2X* is an early event in the development of resistance. To test this hypothesis, we performed NGS of an earlier time point, when cells had largely recovered growth (generation recovery, Gr, in Figure 1B). The results (Figure 3B) showed that indeed chromosome VIII is mostly disomic in all populations already at this time point. Point mutations, instead, were present at very low frequency. The *TUB2* control did not show any relevant disomy at either time point, and mutations of *PHO* genes were entirely missing at the earlier time point (Figure 3B).

The results shown so far support a scenario where the first step in the development of resistance is the acquisition of an extra copy of chromosome VIII, followed by mutations of recurrently mutated genes. At the end of the experiment, most cells are disomic of chromosome VIII and carry one additional mutation. The results suggest that disomy of chromosome VIII and recurrent mutations are adaptive.

### Potential adaptive roles of mutations in recurrently mutated genes

We hypothesized that mutations in the identified genes may be adaptive. Before experimentally testing this hypothesis, we confirmed its plausibility based on the available literature.

*PHO* In *PHO81*, two mutations out of three are nonsense or frameshift, and thus could be classified as loss-of-function. In the case of *PHO4*, four mutations out of five were missense, clustered in the DNA-binding region (Figure S2A). In the hypothesis that they prevent DNA binding, they could also be classified as loss-of-function. Interestingly, uptake of inorganic phosphate is limiting for growth in the cold [34], and deletion of either *PHO81* or *PHO4* rescues growth in low phosphate after deletion of the high-affinity transporter *PHO84* [35].

*KIP3* Among the mutations found in *KIP3*, half are either non-sense or frameshift, again interpretable as loss-of-function. The fact that deletion of *KIP3* reduces sensitivity to benomyl [36] further supports this interpretation, and is in line with the depolymerizing activity of Kip3 [8]. It also reinforces the notion that we selected for mutations that aspecifically contrast induced microtubule depolymerization, and not necessarily the effect of the *tub2-401* allele. The missense mutations found in *KIP3* may underlie subtler mechanisms to recover microtubule polymerization. Recent data point at the L11 domain in Kip3 as essential for microtubule depolymerization [8]. Interestingly, one point-mutation leading to an amino acid change (E359K) was found independently in two evolved populations, and it falls in the L11 domain (Figure 2B). We hypothesize that this mutation disrupts the depolymerase activity of Kip3, while preserving its kinesin function.

*gamma-TuSC* Mutations in *TUB4* and *SPC98* can be understood in the light of increased tubulin nucleation as a means to counteract the *tub2-401* mutations. The amino acid changes in Spc98 are in residues involved in the interaction of Spc98 with Spc97 (P248T) and with Spc110 (H222P), which bridges gamma-TuSC with the centrosome (Figure 4A). Thus, these mutations may stabilize the formation of the gamma-TuSC complex, and favor nucleation. The residue mutated twice in Tub4 (Q12) has been described as important for interacting with GTP, and thus affecting nucleation of microtubules and mediating sensitivity to benomyl [6].

**Figure 4.**
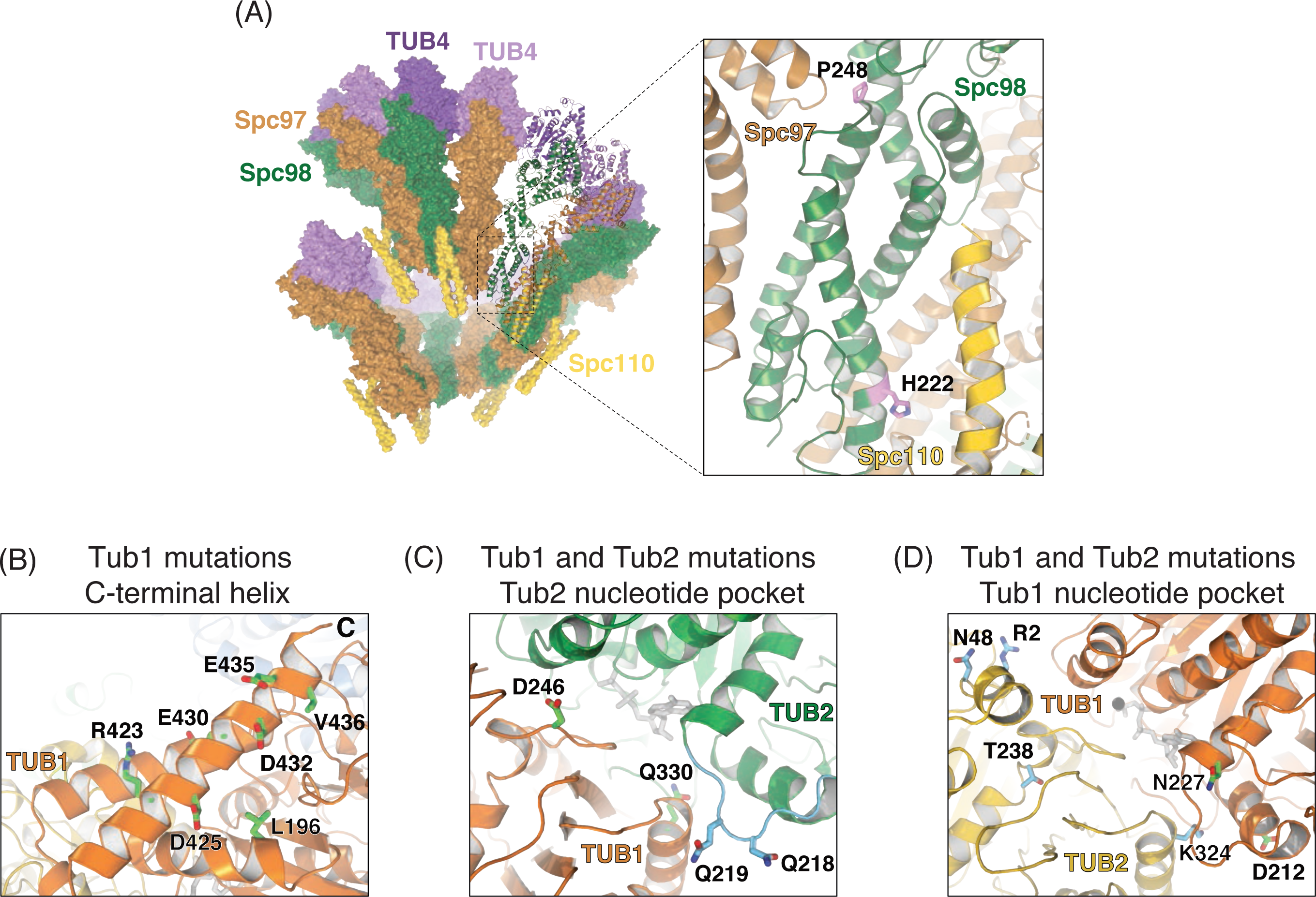
Recurrent amino acid changes in Spc98, Tub1 and Tub2. a) Spc98 residues P248 and H222 are shown in the context of the gamma-TuSC complex (pdb entry 5flz). b) Mutated residues on or near the Tub1 C-terminal helix. c) and d) show residues mutated close to the Tub2 and Tub1 nucleotide pocket, respectively (pdb entry 5W3F).

*TUB1* Among the 12 mutations identified in *TUB1*, we noticed that a large fraction is located on the C-terminal helix 12 (Figure 4B), a region important for interactions with microtubules binding proteins [37]. Hence, we hypothesize that these mutations may alter the dynamics of interaction with binding partners, ultimately stabilizing microtubules. Along the same line, a mutation that occurs twice, L196W is situated close to the C-terminal tail. Its mutation to tryptophan, a residue with a bulky side chain, might misplace the helix, thus inducing a similar effect of the mutants in the helix itself. One of the recurrent mutations is D246Y, which lies at the interface between the Tub1/Tub2 dimer (Figure 4C). Introduction of the bulky tyrosine residue might impair the dynamics of microtubule stability. In agreement with this interpretation, D246A was reported to change sensitivity to benomyl [10]. The same was observed for Ala substitutions in E435, another site on helix 12 that we find mutated twice (Figure 4B).

*TUB2* The three amino acid substitutions in ancestral *tub2-401* (M233V, Y242C, G245L) are located in *TUB2* (Figure S1A). We have not identified revertants among mutations identified in the evolved strains. However, in one population in position 242 we find one change from cysteine to tryptophan. Given the similarity between the latter and the original tyrosine, we can interpret this mutation as reversion. All other mutations (Figure 4C-4D), instead, are possible gain-of-functions. Mutation of T238 into alanine (Figure 4D) has been shown to decrease sensitivity to benomyl [10], with decreasing frequency of catastrophes and slow shrinking [38]. We find it mutated into isoleucine, which can be hypothesized to give a similar phenotype. The change to amino acids with similar chemical properties is confirmed in the recurrently observed N48D mutation (two instances) (Figure 4D), which is located at the Tub2/Tub2 lateral interface and could affect assembly of protofilaments. Similarly, for V229I (two instances) we find an amino acid change that is not expected to change dramatically the chemical properties of beta-tubulin. Yet, it is buried inside the tubulin molecule, and the change may affect its structure. Finally, at the interface between Tub2 and Tub1 we find the most recurrently observed amino acid change, Q219H (6 instances). Q219 belongs to the loop between H6/H7 (highlighted in light blue in Figure 4C), where other mutations have been reported to alter microtubules dynamics [39, 40]. The glutamine to histidine mutation may interfere with tubulin assembly favoring its stability.

In summary, for many of the mutations occurring in recurrently mutated genes, we could provide a rationale for their putative effect on microtubule stabilization based on previous studies pointing at their role for microtubule stability.

### Frequently occurring mutations in recurrently mutated genes are adaptive

To confirm that mutations we identified are adaptive, we introduced some of the most representative ones in the ancestor. In *tub2-401*, we mutated *TUB2* and *KIP3* as each of these genes is altered in 12 out of the 24 populations (Figure 3A, Figure S3B). In the *TUB2* gene (which already carries the mutations of *tub2-401*), we introduced the additional point mutation causing the most frequent amino acid change, Q219H. For brevity, the allele which expresses both the three *tub2-401* amino acid changes and Q219H was called *Q219H*. For *KIP3*, we observed that half of the mutations were nonsense and frame-shifts (Figure 2B). In line with [41, 42], we mimicked such mutations by deleting the gene. We also analyzed the adaptive role of the most widespread disomy, *chrVIII 2X*. As for controls, the most often mutated gene was *PHO4*. Also in this case, we deleted the gene on the assumption that mutations in the DNA-binding site (Figure S2A) impair its transcriptional activity.

We grew cells for 24 hours at 18 °C and measured their growth rate. Both *Q219H* and *kip3D* partially recovered the growth defect of *tub2-401* (Figure 5A-B). Also for *chrVIII 2X* we proved an adaptive role. The effect depended on the *tub2-401* allele, since two copies of the same chromosome impaired growth in the presence of wild type *TUB2* (Figure 5C). Finally, for cells expressing *TUB2*, we observed that growth at low temperature was improved by the deletion of *PHO4* (Figure S4A), in agreement with a role for phosphate uptake in adaptation to the cold [34, 43].

**Figure 5.**
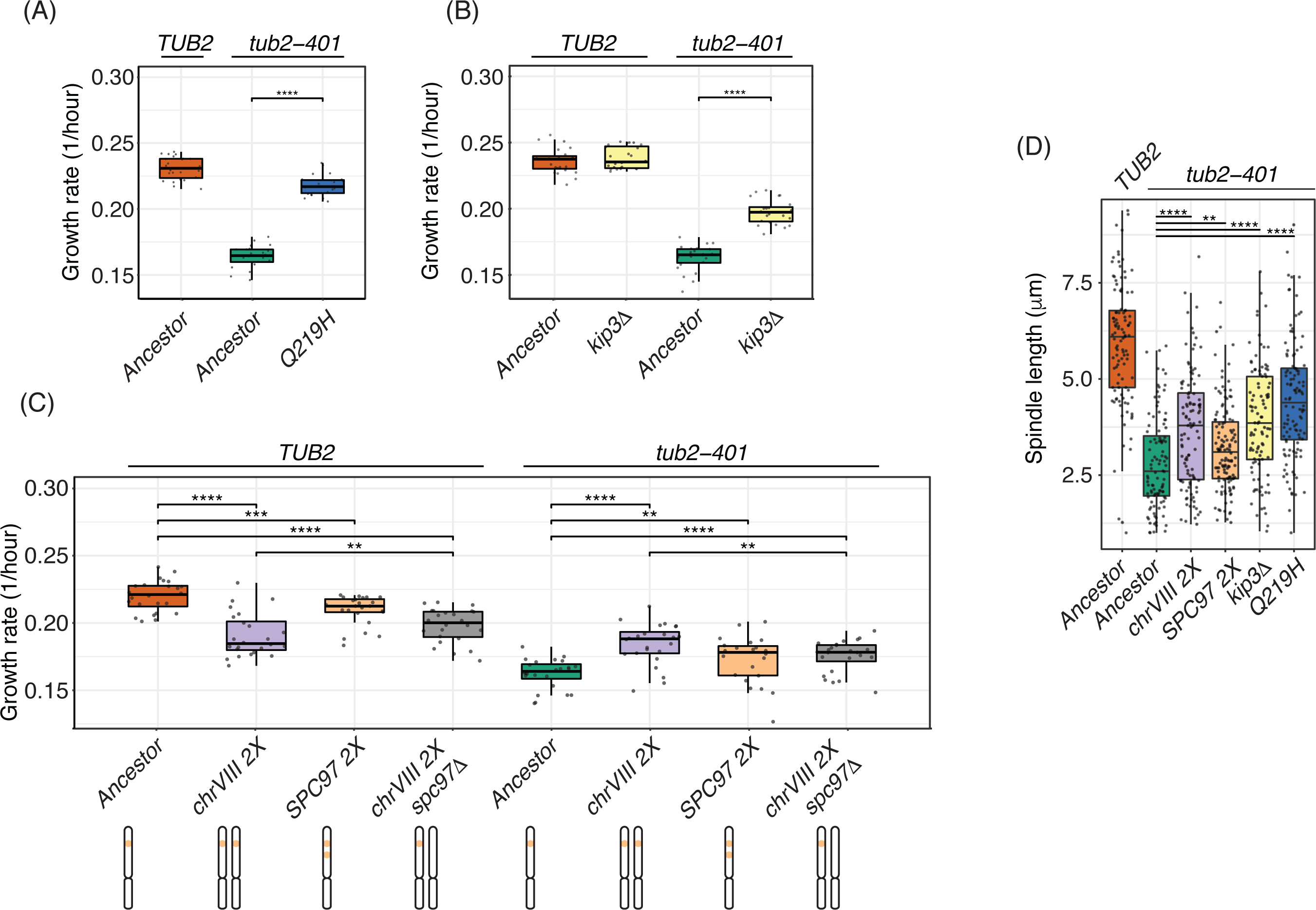
Mutations in recurrently mutated genes are adaptive. a-c) Growth rates measured at 18 °C after growing cells at 18 °C for 24 hours. Pairwise strain comparisons were made using a linear model, adjusting for batch effects for experiments performed on different days. Symbols refer to the p-values of the strain comparison (ns = p-value > 0.5, * = p-value < 0.05, ** = p-value < 10^−2^, *** = p-value < 10^−3^, **** = p-value < 10^−4^). d) Ancestors and mutant strains were synchronized in G1 at 30 °C and released at 18 °C. Tub1 was stained by immunofluorescence while nuclei were stained with DAPI. Spindle lengths were measured with a custom FIJI script across 3 timepoints centered on the maximum fraction of large budded cells (dumbell) (Figure S4B).

We next aimed at identifying whether there is one particular gene on *chrVIII* whose duplication is adaptive. *SPC98*, coding for a component of the gamma-tubulin complex, is mutated in several evolved strains (Figure 2A). Interestingly, another component of the complex, *SPC97*, is located on *chrVIII*. Spc97 is the least expressed among the components of Gamma-TuSC (2.4:1.3:1.0 for Tub4, Spc98, and Spc97) [44]. Hence, we hypothesized that the adaptive effect of the disomy of *chrVIII* may be due to the duplication of *SPC97*. This result is not obvious, since large overexpression of Spc97 was reported to decrease viability [45], and indeed we confirmed a limited reduction of growth due to duplication of *SPC97* in wild type cells (Figure 5C). Accordingly, removing one copy of *SPC97* from the disomic strain carrying *TUB2* wild type improved slightly its growth rate. However, when we introduced two copies of *SPC97* in a strain expressing *tub2-401* and monosomic for *chrVIII*, we observed an improvement of growth rate comparable to that of disomic strains. Importantly, deletion of one copy of *SPC97* in a strain carrying *tub2-401* and disomic for *chrVIII* decreased growth compared to the disomic strain. However, it did not revert it completely to that of *tub2-401* alone (Figure 5C), implying that additional genes located on *chrVIII* contribute to the increased fitness of disomic strains. These results suggest that when microtubule polymerization is crippled, duplication of *SPC97* provides an advantage that overcomes the costs coming with duplication of *chrVIII*.

To confirm that the adaptive effect is due to the increased stability of microtubules, we analyzed mitotic spindles of cells where we engineered recurrent mutations in the presence of *tub2-401*. Cells were synchronized in G1, released at 18 °C and collected at different time points when they were arrested in mitosis (ie, with large buds, Figure S4B). Mitotic spindles were on average longer in cells carrying the recurrent genetic changes when compared to the ancestor *tub2-401* (Figure 5D, Figure S4C).

We conclude that the recurrent mutations we have tested are adaptive. The disomy of chromosome VIII confers resistance to microtubule depolymerization partially due to the duplication of *SPC97*.

### Epistatic interactions

Next, we aimed at understanding the evolutionary trajectories leading to resistance. To this aim, we combined the three genetic changes analyzed so far: *Q219H*, disomy of *chrVIII* and *kip3D*, always in the presence of the *tub2-401* allele (Figure 6A).

**Figure 6.**
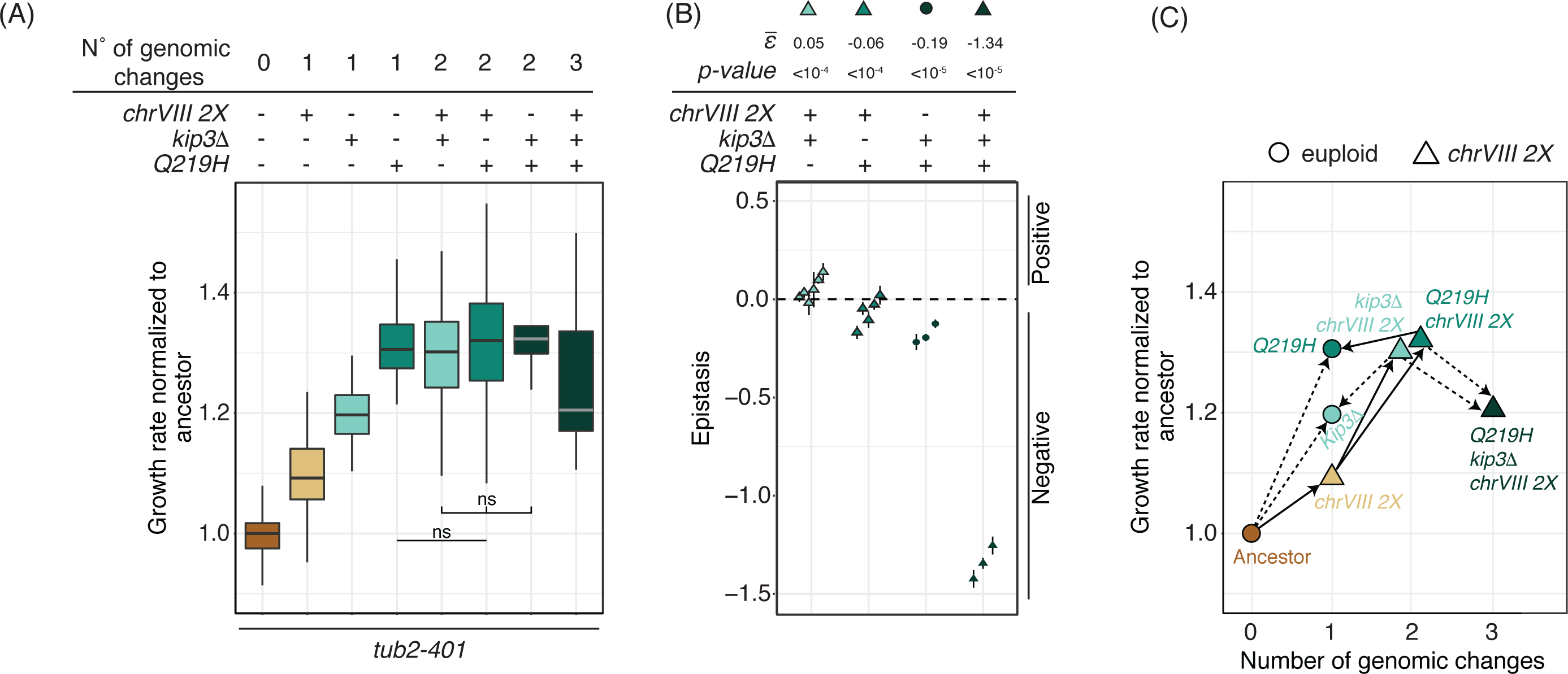
Epistatic effects of adaptive mutations. a) Growth rates measured at 18 °C after growing cells at 18 °C for 24 hours. All growth rates are normalized to the value of the ancestor (*tub2-401*). Pairwise strain comparisons were made using a linear model, adjusting for batch effects for experiments performed on different days. Symbols refer to the p-values of the strain comparison (ns = p-value > 0.5, * = p-value < 0.05, ** = p-value < 10^−2^, *** = p-value < 10^−3^, **** = p-value < 10^−4^). The group comparison among strains with 2 genomic changes is made using Kruskal-Wallis test (p-value = 0.149). b)Epistatic interactions among adaptive mutations. Epistasis is the difference between the observed fitness and the one calculated summing the fitness of individual mutations. Each symbol represents one experiment with 8 replicates for each strain, and bars are the standard error. ε is computed as the weighted mean over independent experiments. The calculations were done assuming normalized growth rate as a proxy for fitness. See Material and methods for details. c) The median values of the measurements shown in (a). On the x-axis, we report the number of genetic changes, i.e. point-mutations or changes in ploidy. Solid lines identify changes that we propose to have occurred during emergence of resistance. Dotted lines, instead, are changes that have not occurred. For sake of clarity, *Q219H kip3Δ* is excluded from this panel.

*Q219H* was the most effective single mutation in terms of increase of growth rate, followed by deletion of *KIP3* and disomy of *chrVIII*. Any pair-wise combination of these three gave a similar increase in growth rate over the ancestor *tub2-401*. Hence, additional mutations had the least effect on *Q219H*, which alone showed a behavior most similar to the wild type (Figure 5A). With the assumption that growth rate is an approximation of fitness, we could quantitatively compute the epistatic interactions. Indeed, we found that *Q219H* shows negative epistasis with either *chrVIII* 2X or *kip3Δ* (Figure 6B). Both *kip3D* and *chrVIII 2X*, instead, starting from a lower growth rate benefited from the addition of any of the other two genetic changes (compare column 3, 5 and 7 for *kip3D* and 2, 5 and 6 for *chrVIII 2X* in Figure 6A). Their interaction showed a slight positive epistasis (Figure 6B). Finally, when we introduced in the same strain all three changes, we observed that the triple mutant fared worse than double mutants (Figure 6A), showing the strongest negative epistasis (Figure 6B). This result further supports the interpretation that evolved strains at the final generation Gf carry disomy of *chrVIII* and only one additional mutation in the genes we identified.

We then drew the fitness landscape and analyzed the evolutionary trajectories that cells have taken to develop resistance (Figure 6C). The analysis is useful since it allows to compare actual trajectories to all those potentially possible. It is worth noticing that cells seem to have avoided the most direct evolutionary trajectory to *Q219H*, the fittest solution together with *Q219H chrVIII 2X* (Figure 6A). We already noticed that at the time of recovery (Gr) (Figure 3B) cells show widespread disomy of *chrVIII* and very limited point-mutations. Hence, possibly due to the increased chromosome missegregation rate (Figure S1B), the first step along the emergence of resistance is disomy of *chrVIII*. Only later, cells acquired point mutations. Some of these, like deletion of *KIP3*, increase their fitness when combined with *chrVIII 2X*, and thus these cells are unlikely to become euploid. Others, like *Q219H*, have the same fitness with or without disomy of chromosome VIII, and may find advantageous becoming euploid again. Accordingly, we only observe a decrease of aneuploidy in the final generation Gf within populations that have mutations in *TUB2* (Figure 3A).

We conclude that adaptive genetic changes leading to resistance to impaired microtubule polymerization display epistatic interactions. Mutations in *TUB2* give the largest fitness increase. Nevertheless, they are not the first to occur but follow disomy of *chrVIII*. The same is true for the other point mutations. Whether cells will eventually keep the mutations and loose the disomy of chromosome VIII will likely depend on the fitness gain coming with the adaptive mutation alone.

## DISCUSSION

We evolved cells expressing mutations impairing microtubules polymerization when grown at low temperature. The mutations cause a large increase in chromosome missegregation. After roughly 150 generations, evolved cells were again capable to assemble proper mitotic spindles. We evolved in parallel 24 different populations. Remarkably, they were characterized by a simple karyotype with only one disomic chromosome (*chrVIII*) across most populations. The pattern of recurrently mutated genes is also simple, and includes only two classes: tubulins and kinesins. By introducing in the ancestors strains the disomy or the most frequently observed mutations in these two classes, we could ascertain their adaptive nature and analyze their epistatic interactions.

The two adaptive processes (change of ploidy and point mutations) are not interchangeable, but follow a precise temporal sequence. When we analyzed populations that had partially recovered growth at an earlier time point, we found that most of them carried disomy of chromosome VIII with high frequency (>80%). On the contrary, there was a paucity of single nucleotide mutations. Hence, we propose that disomy of chromosome VIII provides the first compensatory solution against the impairing mutations. We further showed that this was at least partly due to the duplication of *SPC97*, a member of gamma-TuSC. At a later time-point, sequencing confirmed the presence of disomy for chromosome VIII in the large majority of populations. In addition, cells in all populations had mutated one, but not more, of the recurrently affected genes.

Hence, disomy of chromosome VIII comes first, mutations of tubulins or *KIP3* follow. The repetition of these evolutionary paths of compensatory mutations in several independent experiments shows that the emergence of resistance is potentially predictable.

### Aneuploidy is an adaptive solution to impaired microtubule polymerization

The role of aneuploidy as a fast adaptive solution has been reported for several sources of stress [22, 25, 26, 46]. Indeed, stimuli that increase chromosome missegregation rate generate genetic variability which favours the arising of resistance to other treatments as well [23]. Given that impaired microtubule polymerization increases missegregation rate, it is not surprising that the first compensatory event we observed was disomy. Whether this solution persists in time, instead, is not obvious. Aneuploidy is a fast but costly solution. As such, in other contexts it was shown to be replaced on the long run by more precise and less detrimental point mutations [25]. However, our data show that this is not necessarily the case. For example, in the case of *KIP3* deletion the combination with an extra copy of chromosome VIII does not show a detrimental effect. In this case, we do not expect a loss of disomy of chromosome VIII as long as Kip3 is dysfunctional. On the contrary, for the tubulin mutations *Q219H*, the loss of extra chromosome VIII is not a disadvantage. Accordingly, we observed a few populations that carry only mutations in *TUB2* at the end of the experiment, while all were disomic at an earlier time point. In brief, the persistence of aneuploidy as an adaptive solution will depend on the epistatic interaction with the additional compensatory mutations.

Recently, a study analyzed the effect of increasing chromosome missegregation by different means, ie inactivation of the chromosome passenger complex via deletion of the survivin homolog *BIR1* [24]. The adaptive solutions they found are partially different from ours. There are no recurrently mutated genes, and among the genes mutated there are not those that we identify as recurrently mutated. This is maybe not surprising, since microtubule polymerization is not affected by the deletion of *BIR1*. Aneuploidy, instead, is a major adaptive event in both experiments. The evolved karyotypes partially overlap, but are not identical. Both studies identify disomies in *chrVIII* and *chrIII* (the latter being very limited in our study). Ravichandran, Campbell, and co-workers, however, identified more complex karyotypes which include disomies of *chrII* and *chrX*. Hence, how cells react to chromosomal instability at least in part depends on the stimulus that triggers it. However, it will be important to understand whether specific aneuploidies provide a generic mechanism to cope with high chromosome missegregation rates *per se*.

### Compensatory mutations are largely due to gain-of-functions

Both high-throughput studies and specific in-depth analyses in yeast have shown that cells can adopt different strategies to recover from growth defects: aneuploidies, gain-of-function mutations, loss-of-function mutations, large structural variants [27]. However, not all strategies have the same probability to occur. Previous evolution repair experiments indicated that loss-of-function mutations tend to occur more often than gain-of-functions [21]. In our case, we observed quite the opposite. Multiple gain-of-function mutations occur in alpha-, beta-, and gamma-tubulins, as well as in the other elements of the gamma-TuSC complex. Only in *KIP3* we found loss-of-function mutations, ie, nonsense or frameshift mutations. In this gene, we also identified several missense mutations which can also be interpreted as loss-of-function, likely partly preserving enzymatic activity (eg, the two mutations occurring in the L11 domain).

Why so many gain-of-function in our evolution experiment? Compared to previous studies [22, 41, 42], an obvious difference is that we did not start our experiment with a deletion but with 3 impairing amino acid substitutions. As such, the gain-of-function mutations are in fact recovery-of-function. With this term, we do not mean revertants, which we observe only in one case. Rather, we refer to mutations that can make use of the mutated and partially dysfunctional tubulin. Such mutations can be intragenic, occurring in *TUB2*, but many of them affect other genes whose products interact directly with beta-tubulin.

### What can we learn from laboratory evolution experiments on the mechanisms leading to resistance to antimitotics?

Tubulins, and also their main interactors such as kinesins, are conserved among eukaryotes. Adaptive mutations observed in our strains affect genes that are also involved in the development of resistance in human cells. Kinesins play a role in the resistance to drugs affecting microtubule dynamics [18, 19]. 50% of the residues that we found mutated in *TUB2* align with residues that were identified as mutated in a recent study in cancer patients [16]. Of course, mammals have additional complexity and therefore additional routes to resistance such as differential expression levels of the many different tubulin isotypes. Nevertheless, we propose that our results may point to general principles in the emergence of resistance that may also apply to mammalian cells. It is worth noticing that these results could not be obtained simply by developing and analyzing resistant cell lines in the lab. Our experiment offered the unique opportunity to follow the emergence of mutations in a time-resolved manner.

First, since antimitotics are bound to increase chromosome missegregation, we propose that aneuploidy plays a key role in the early steps of the development of resistance also in mammals. Our data suggest that this is the case even if aneuploidy does not provide the best adaptive advantage. We suspect that aneuploidy as a cause of resistance has gone unnoticed so far either because studies have focused on established resistant cells (when aneuploidy may have been lost), or simply because it has not been searched for. Second, the high reproducibility of the path leading to resistance supports the concept of preventing or delaying resistance by targeting specific few and well-defined molecules during the proper time window. Third, different adaptive mutations may display different epistatic interactions, like those we observed between *chrVIII 2X* and either *Q219H* or *kip3D*. This point and the previous one have potentially relevant implications. Interfering with the development of resistance one may paradoxically select cells that are more efficiently resistant (eg, contrasting disomy of *chrVIII* may select *Q219H*, which may turn out to be even fitter than *Q219H chrVIII 2X*). This result calls for the importance of understanding epistatic interactions for devising efficient strategies aimed at contrasting the development of resistant cells. In conclusion, we showed that yeast cells can be effectively used to rationalize complex evolutionary processes taking place in higher eukaryotes.

## ACKNOWLEDGMENTS

We thank Zoltan Farkas for introducing us to fitness measurements. We thank Zoltan Farkas, Silke Hauf, Claudio Vernieri and Andrea Musacchio for constructive comments on the manuscript. Work in the group of AC is financed by AIRC, the Italian association for cancer research (Grant AIRC-IG 21556); MP benefits of a AIRC fellowship; SeP receives supports from the Italian Ministry of Health (Ricerca Corrente and 5×1000 funds); SiP benefits from a Fondazione Umberto Veronesi fellowship; MCL is supported by AIRC, the Italian association for cancer research (Grant AIRC-IG 23258); DS is supported by the National Research, Development and Innovation Fund of Hungary (grants K_124881 and FIEK_16-1-2016-0005); GR is supported by a Singapore NRF Investigatorship (NRF-NRFI05-2019-0008); the Hungarian Academy of Sciences sponsored the collaboration between the groups of AC and DS by awarding a visiting fellowship to AC.

---

**Figure S1 in support of Figure 1.**
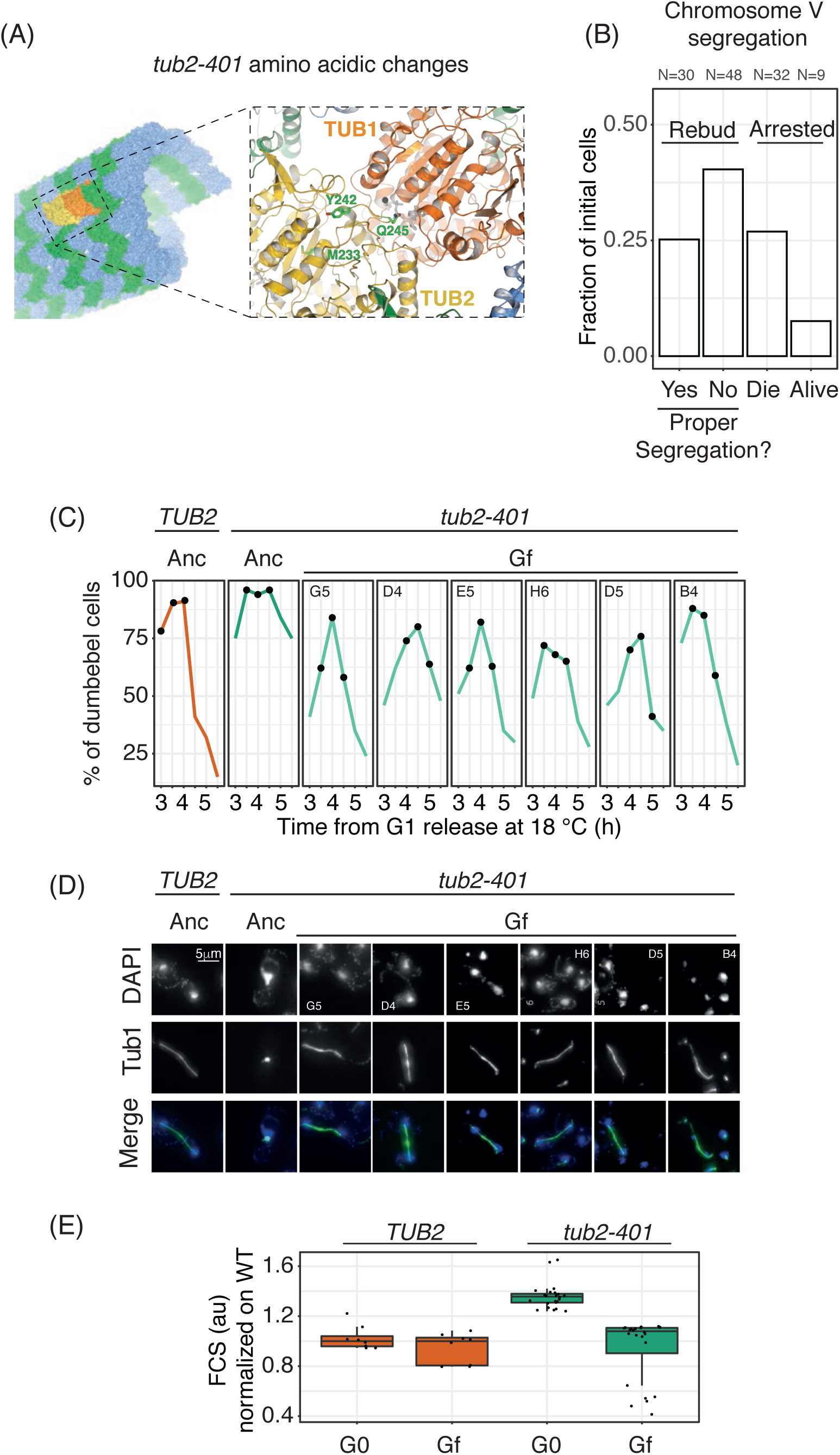
Characterization of evolved cells. a) The three amino acid changes of the *tub2-401* allele. b) Percentage of missegregating cells in the ancestor *tub2-401*. Cells were synchronized in G1 at 30 °C, the temperature was shifted to 18 °C during the last hour of synchronization, and finally cells were released from the arrest at 18 °C in the microfluidic chamber. The number of missegregating events was measured by following the first cell division. The events of missegregation were identified by following GFP-tagged chromosome V. c) Cells sampled in the last time point (Gf) and ancestors were synchronized in G1 at 30 °C and released at 18 °C. The fraction of cells with a large bud was monitored every 30 min from the G1 release. Black dots identify the timepoints in which the spindle lengths were measured (Figure 1C). d) Representative spindles in Ancestor and Gf cells (Figure 1C). Tub1 was stained for immunofluorescence while nuclei were stained with DAPI. e) As a proxy for cell size, we used forward scattering measured at FACS (Macsquant). G0 cells were fixed after 24 hours at 18 °C (first timepoint of evolution experiment) while Gf cells were collected at the end of the experiment (Figure 1B).

**Figure S2 in support of Figure 2.**
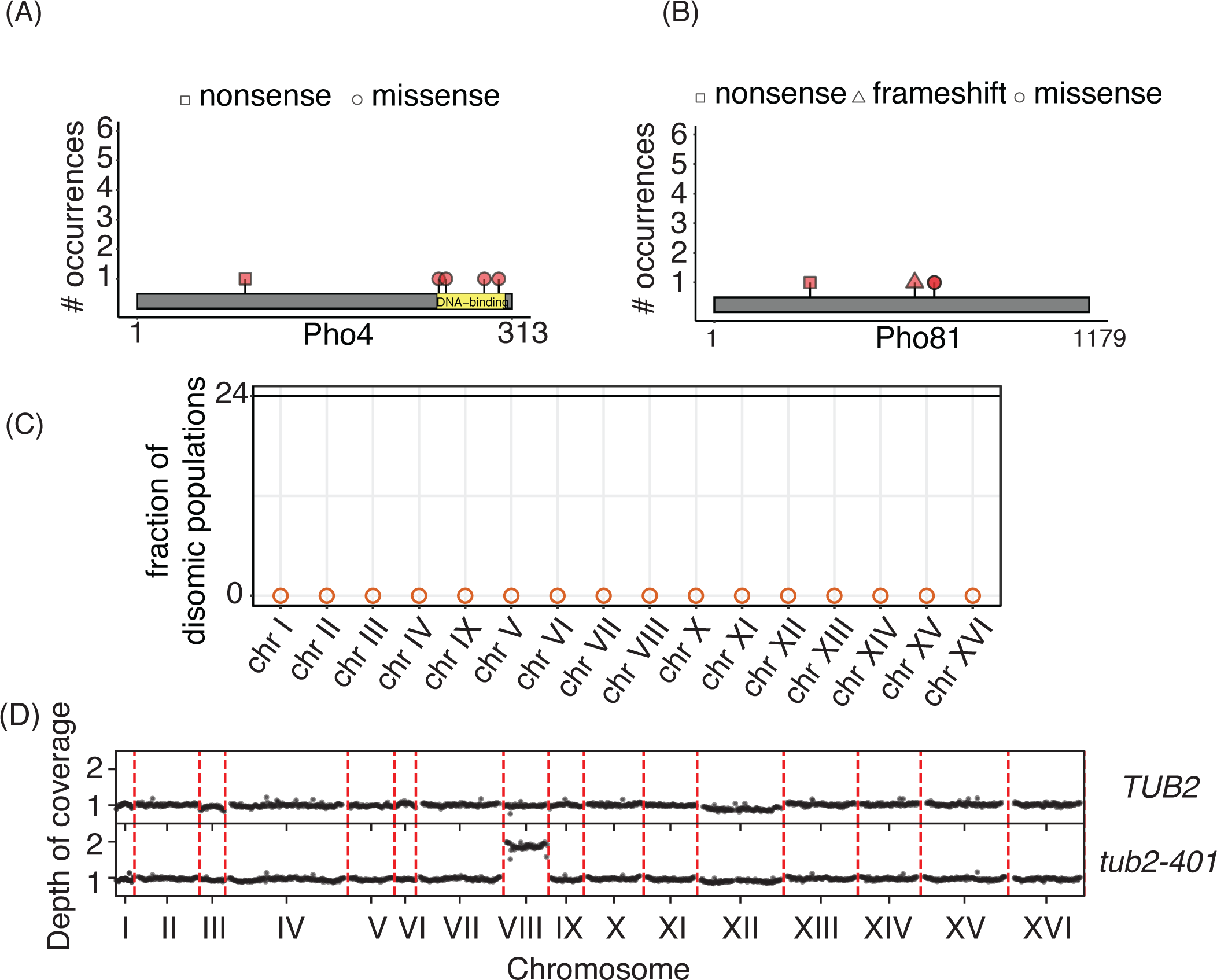
Recurrent amino acid changes in the PHO pathway and disomies, at the end of the experiment. a)-b) Amino acid changes caused by mutations occurring independently multiple times in *PHO81* and *PHO4*. They were detected only in *TUB2* cells (Figure 3) c) Fraction of disomic populations. Empty dots are used when there are no disomic chromosomes. Chromosome copy numbers were determined by coverage analysis (an example in Figure S2D). d) Normalized and corrected depth of coverage for two representative samples (*TUB2*, population A1; *tub2-401*, population A4). Each dot represents the median depth over a 10.000 bp window.

**Figure S3 in support of Figure 3.**
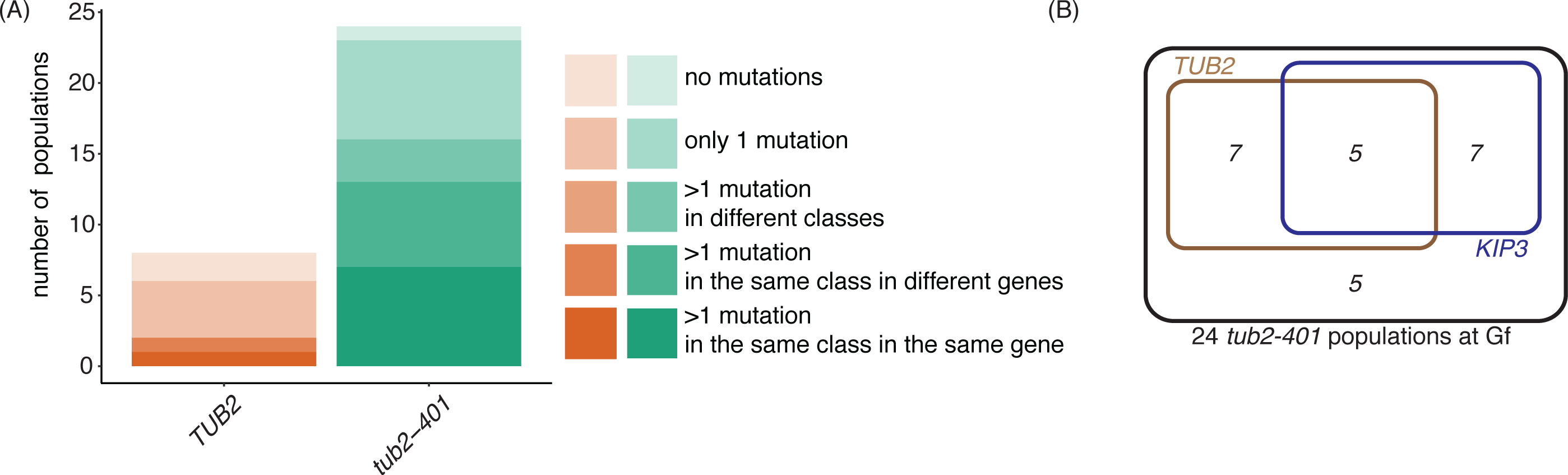
Mutation frequencies at final timepoint (Gf) a) Bar plot of populations grouped according to the presence of multiple mutations in recurrently mutated genes. b) Venn Diagram of *tub2-401* at Gf, carrying mutations in *TUB2* or *KIP3* (data shown in the third panel from top of Figure 3A).

**Figure S4 in support of Figure 5.**
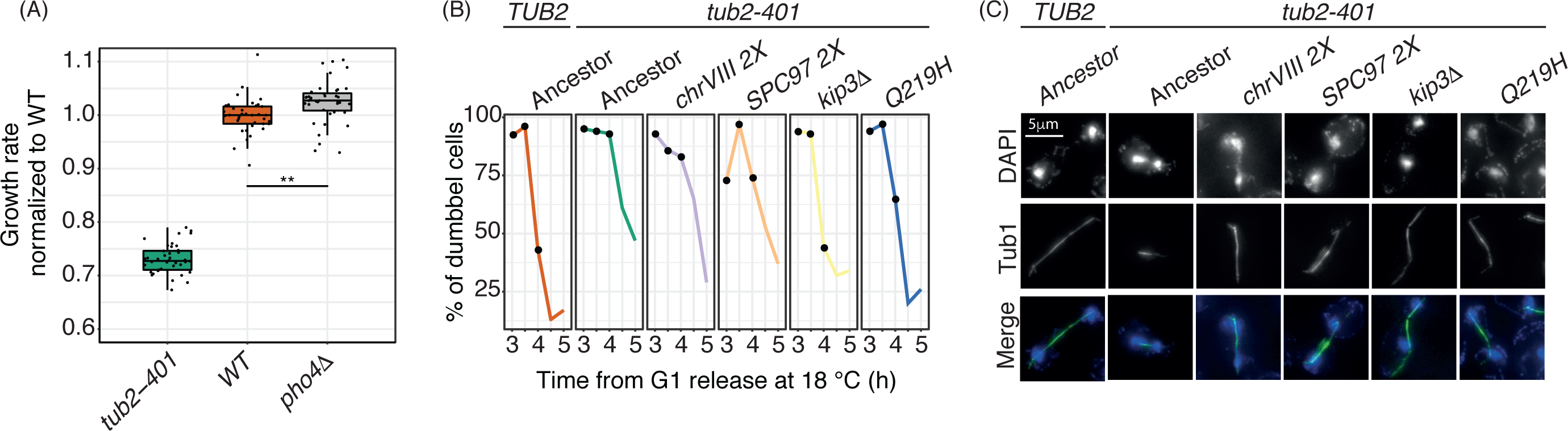
Characterization of engineered strains. a) Growth rates of *TUB2, tub2-401* and *TUB2 pho4D* were measured at 18 °C across 8-hours after growing cells at 18 °C for 24 hours. b) - c) Spindles measurements in *tub2-401* cells carrying adaptive mutations. Cells were synchronized in G1 at 30 °C and released at 18 °C. b) The fraction of large-budded cells was monitored every 30 minutes from G1 release. Black dots identify the timepoints in which the spindle lengths were measured (Figure 5D). c) Representative spindles. Microtubules were identified by immunofluorescence on Tub1 while nuclei were stained with DAPI.

**Table S1.**
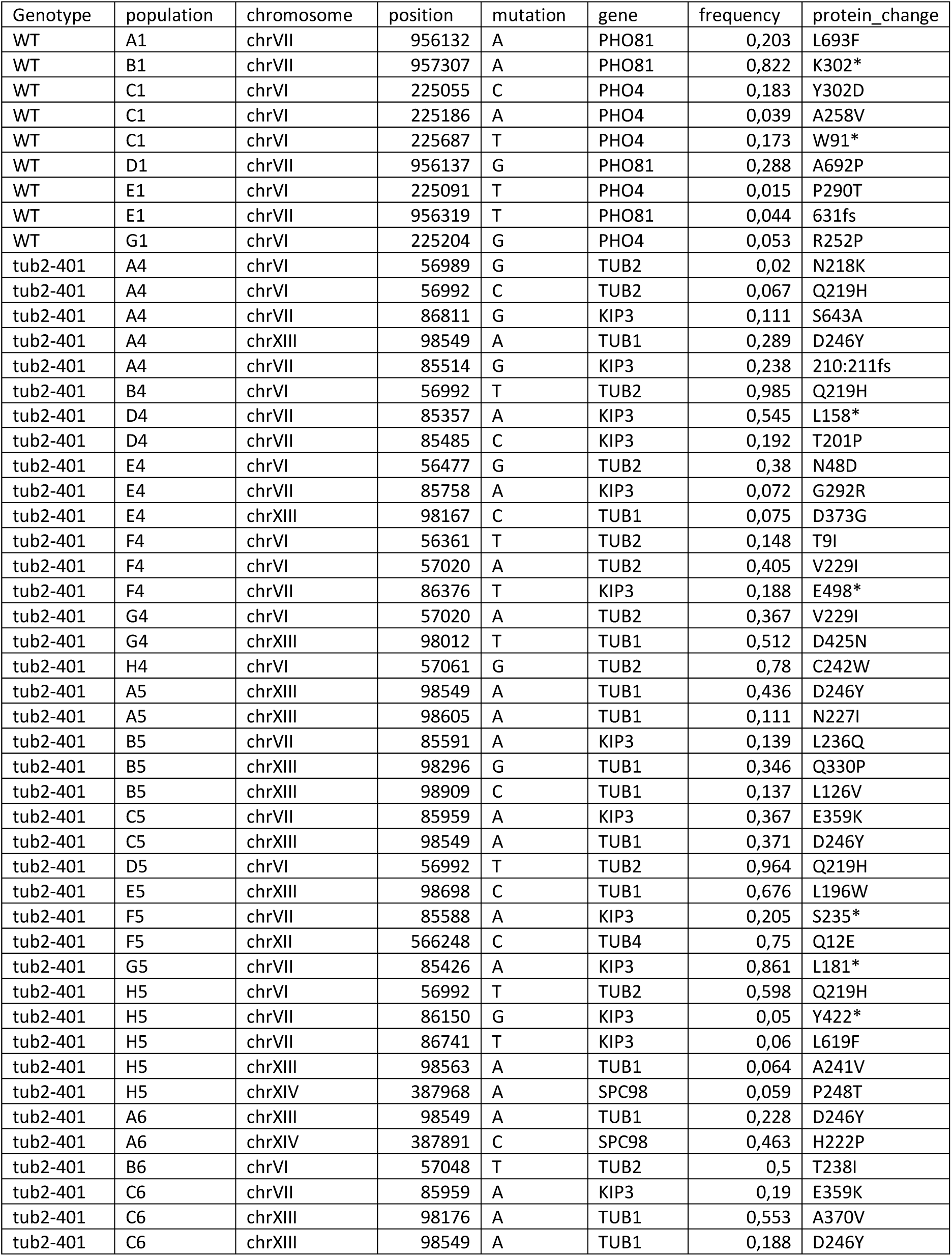
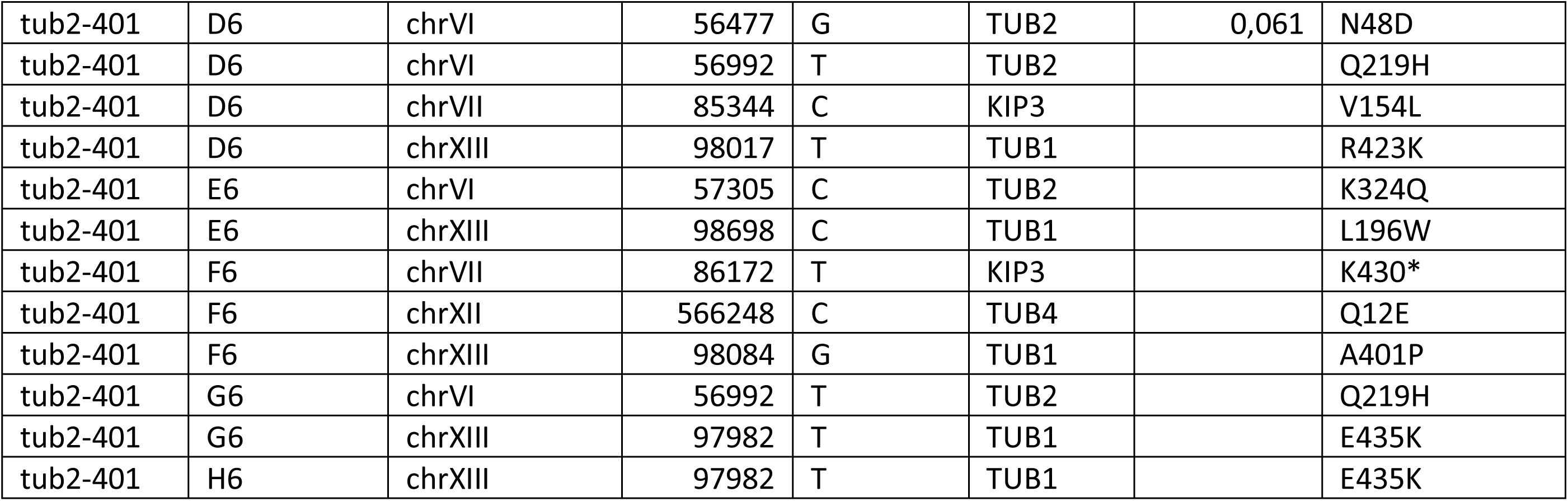
List of recurrently mutated genes. The list includes all genes that are mutated more than once, with different mutations, in at least two populations. See Material and Methods for details.

**Table S2.** Adaptive mutations identified in clones derived from mixed populations. Clones were derived from *tub2-401* populations at the end of the experiment (Gf). We derived them from populations where multiple adaptive mutations have high frequencies (see Figure 3). For each clone, we tested by Sanger sequencing the presence of individual mutations.

|  |  | *Number of clones with mutations in the indicated genes* |  |  |  |  |
| --- | --- | --- | --- | --- | --- | --- |
| **Populations** | **Recurrently mutated genes** | ***TUB1*** | ***TUB2*** | ***TUB4*** | ***KIP3*** | **Concurrent mutations** |
| C5 | *TUB1, KIP3* | 4 | - | - | 2 | 0 |
| F4 | *TUB2, KIP3* | - | 6 | - | 1 | 0 |
| F5 | *TUB4, KIP3* | - | - | 8 | 0 | 0 |
| G4 | *TUB1, TUB2* | 3 | 4 | - | - | 0 |

**Table S3.** List of strains. All strains used in this study are derivatives of a modified W303 prototroph for *URA3* and *TRP1* (*leu2-3, can1-100, ade2-1, his3-11,15 ura:URA3(1X), trp1:TRP1(2X)*)

|  |  |
| --- | --- |
| yAC4248 | *MATa*, *pho4::KanMX6* |
| yAC4256 | *MATa*, *kip3::KanMX6* |
| yAC4258 | *MATa*, *kip3::KanMX6*, *tub2-401* |
| yAC4314 | *MATa*, haploid disomic chr VIII, *DOG2-HA3::HPH*, *DOG2-myc9::NAT*, *tub2-401* |
| yAC4355 | *MATa*, *spc97:SPC97(2x):NAT* |
| yAC4357 | *MATa*, *spc97:SPC97 (2x):NAT*, *tub2-401* |
| yAC4371 | *MATa*, haploid disomic chr VIII, *DOG2-HA3::HPH*, *DOG2-myc9::NAT*, *kip3::KanMX6*, *tub2-401* |
| yAC4376 | *MATa*, *tub2-401A657T* |
| yAC4383 | *MATa*, haploid disomic chr VIII, *DOG2-HA3::HPH*, *DOG2-myc9::NAT* |
| yAC4384 | *MATa*, haploid disomic chr VIII, *DOG2-HA3::HPH*, *DOG2-myc9::NAT*, *tub2-401A657T* |
| yAC4397 | *MATa* |
| yAC4398 | *MATa*, *tub2-401* |
| yAC4399 | *MATa*, haploid disomic chr VIII, *DOG2-HA3::HPH*, *DOG2-myc9::NAT*, *spc97::KanMX6* |
| yAC4400 | *MATalpha*, haploid disomic chr VIII, *DOG2-HA3::HPH*, *DOG2-myc9::NAT*, *spc97::KanMX6*, *tub2-401* |
| yAC4472 | *MATa*, haploid disomic chr VIII, *DOG2-HA3::HPH*, *DOG2-myc9::NAT*, *tub2-401A657T*, *kip3::KanMX6* |

**Table S4.** List of plasmids.

| **number** | **description** | **origin** |
| --- | --- | --- |
| pAC67 | pFA6a-kanMX6 | [1] |
| pAC146 | pFA6a-3myeGFPdcufc:natNT1 | gift from Michael Knop lab |
| pAC149 | tub2-401 in Yip5 | [2] |
| pAC173 | tub2-401(Q219H) in Yip5 | This study |

**Table S5.**
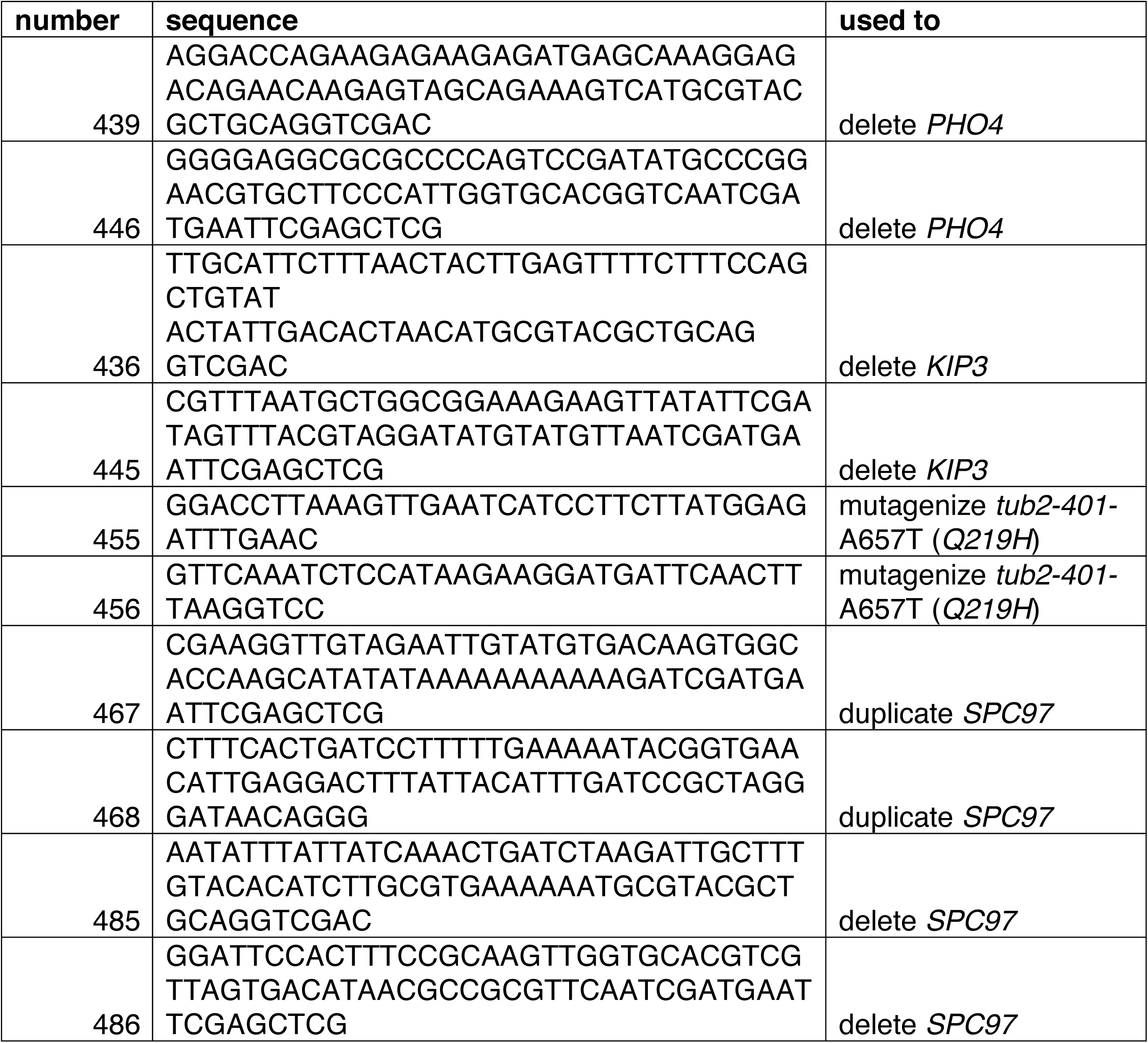
List of oligos.

**Table S6.** List of unique mutations in ancestors *tub2-401* and *TUB2*.

| *Genotype* | *chromosome* | *position* | *mutation* | *gene* | *protein_change* |
| --- | --- | --- | --- | --- | --- |
| WT | chrIV | 1428232 | A | intergenic | NA |
| WT | chrX | 544304 | C | CDC8 | K81N |
| tub2-401 | chrIV | 389636 | G | GPR1 | S808P |
| tub2-401 | chrVII | 1049338 | T | SCW4 | NA |
| tub2-401 | chrX | 165092 | T | AIM23 | R86K |
| tub2-401 | chrXI | 46108 | A | STE6 | S63L |
| tub2-401 | chrXIV | 780662 | T | COS10 | L249F |
| tub2-401 | chrXII | 818842 | A | intergenic | NA |
| tub2-401 | chrXII | 1013798 | GT | intergenic | NA |

## Yeast strains

All yeast strains listed in Table S3 are derivatives of or were backcrossed at least five times with W303. Here, we used a modified W303, protrotroph for uracil and tryptophan (*ura3::URA3* and *trp1::TRP1 (2X)*). The evolutionary experiment was performed using MATa haploid strains.

Deletions were obtained by one-step gene replacement, following S-primer strategy described in [1]. For *PHO4, KIP3*, and *SPC97* deletions, PCR was made on plasmid pl67 (pFA6a-kanMX6 [2]) to obtain KanMX cassette. *SPC97* deletion was performed on *chrVIII* disomic strains, since the gene is essential.

To obtain *tub2-401*^*A657T*^ mutants (*Q219H*), plasmid pAC149 carrying *tub2-401* allele (a gift from T. Huffaker, Department of Molecular Biology and Genetics, Cornell University - Ithaca, United States) was mutagenized by PCR using primers 455/456 to obtain plasmid pAC173. Then, following [3], this plasmid was digested with KpnI to transform yeast, directing the integration at *TUB2* locus. Transformants were plated on 5-fluoro-orotic acid to select clones that excided the *URA3* marker, and the resulting colonies were checked by drop test at 18 °C and compared with *tub2-401* mutants: the ones that grew better than *tub2-401* were checked by Sanger sequencing to confirm the presence of the *tub2-401*^*A657T*^ allele.

To duplicate *SPC97*, we followed [4]. We duplicated the genomic region from the end of *ARS820C* (first gene upstream *SPC97*, on the other strand) to the first codon of *YHR173C* (first gene downstream *SPC97*, on the other strand). In this way, we kept 5’- and 3’UTRs of *SPC97*, avoiding duplicating the *ATG* of *YHR173C*.

Plasmids and oligos described above are listed in Tables S4 and S5

## Media and growth conditions

Cells were grown in YP medium (1% yeast extract, 2% Bacto Peptone, 50 mg/l adenine) supplemented with 2% glucose (YPD). For the evolutionary experiment, YPD medium was supplemented with Penicillin-Streptomycin 100X (BioWest).

To arrest cells in G1, α-factor [5 μg/ml] was added to cells growing exponentially at 30 °C. After 1 hour and 30 minutes, α-factor was re-added at half the concentration [2.5 μg/ml]. If cells have to be released from G1 arrest at 18 °C, after 2 hours from the first α-factor addition cells were released in fresh medium at 18 °C supplemented with α-factor [5 μg/ml] and left grown at 18 °C for 1 hour before release from the G1 arrest.

## Evolution experiment

Ancestral strains were thawed and isolated on YPD plates at 30 °C. 24 independent colonies of *tub2-401* and 8 of *TUB2* were inoculated at 30 °C in 800 μl of YPD in 96-deepwell plates and shifted at 18 °C the day after. During cell growth, plates were covered with aeraseals (Sigma) and incubated at 18 °C under constant orbital shaking on deepwell block tilted adaptors. All the clones were cultured in the same 96-deepwell plate and handled with a TECAN Freedom EVO 150 liquid handler. Cells were diluted twice a day in fresh cold YPD (18 °C) to maintain the exponential phase, and assessed for growth rate every 3-4 days.

To measure growth rate, cells were diluted at OD_600_ 0.2 in 96-microtiter plates (NUNC - Lifetechologies) and cultured at 18 °C for 8 hours while measured every hour using a Cytation 5 (BioTek) OD_600_ plate reader. For each timepoint, OD_600_ of each well was acquired 5 times, and the mean value was used for further analysis.

For each experiment, one or more wells were filled with clean medium, both to check for contamination and to have a measure of background OD_600_. The background OD_600_ was subtracted to the raw OD_600_ to compute the net OD_600_. For each well, the net OD_600_ was fitted against the actual time of measurement to estimate apparent growth rate (α), following the equation:

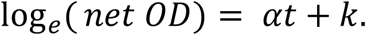

To control for ‘goodness of fit’, we used Root-mean-square deviation RMSD, defined as:

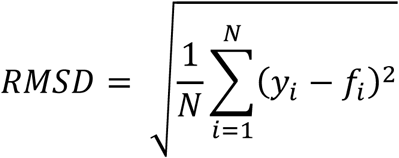

where *N* is the number of measurements, *y*_*i*_ are the measured values, and *f*_*i*_ are the fitted values.

Wells with RMSD greater than 0.1 were excluded from the results.

### Growth Assay (GA) on Engineered strains

Cells were inoculated in glass flask in YPD medium at 30 °C and grown overnight. The day after, each culture was diluted at OD_600_ 0.1 and divided in 8 wells of a deepwell block plate (Greiner-Bio One) and grown for 24 hours at 18 °C under constant shaking. Then, cells were diluted in 96-well microtiter plates (NUNC - Lifetechnologies) at net OD_600_ 0.025 in 200 μl of fresh and cold (18 °C) YPD medium and incubated at 18 °C under constant shaking. OD_600_ were measured every hour for 8 hours using TECAN 200M Infinite plate reader. For each timepoint, OD_600_ of each well was acquired 5 times, and the mean value was used for further analysis. Measurements with OD_600_ above 1.5 or below 0.01 were removed, since the linearity range sits between these values (data not shown).

Growth rate was measured as presented above for the evolution experiment.

### Epistasis

Epistasis ε is the difference between the measured and the expected fitness of the multiple mutant assuming that the mutations are independent. Following [5] (equation 26), expected fitness 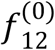 is computed as

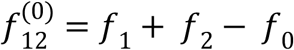

where *f*_1_ and *f* _2_ are the fitness of the two mutants, and *f* _0_ is the one of the ancestor. By defining *f* _12_ as the measured fitness of the multiple mutant, epistasis is computed as

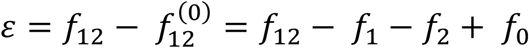

For each experiment, <: is the median of the normalized growth rate of 8 technical replicates of the *i-*th strain. Standard error of the epistasis is computed as the root of the sum of the squared standard errors of each *f*1

The final epistasis 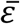 is the weighted average of the replicates, using the standard errors as weights. Standard error of the weighted mean is

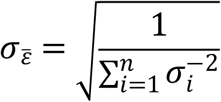

The *p*-value is a standard one-tail *p*-value test (right tail for positive epistasis, left tail for negative epistasis) where we have used a gaussian approximation with mean 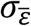 and standard deviation FKE.

### Tubulin immunofluorescence

Cells were arrested in G1 as described above, and released at 18 °C. Samples were collected every 30 minutes from 3 to 5 hours after the release. For each strain, the 3 timepoints with the highest fraction of large-budded cells (dumbbell) were treated for immunofluorescence as follows. 1 ml of cells was collected and fixed with 3.7% formaldehyde in KPi buffer (0.1M Kphos pH6.4 0.5mM MgCl_2_) and incubated overnight at 4 °C.

Cells were washed 3 times with KPi buffer and once with Sorbitol solution (1.2 M Sorbitol, 0.1 M KPi pH 7.4, 0.5 mM MgCl_2_). Then, cell wall was digested at 37 °C with 200 μl of Sorbitol solution supplemented with 5 μl of zymolyase 10 mg/ml and 2 μl of 2-mercaptoethanol for 15/20 minutes and washed with Sorbitol solution. 5 μl of spheroplasts were loaded to glass slides (Thermo-Scientific) coated with polylysine (Sigma-Aldrich). After 15/20 minutes, slides were immersed in cold MeOH (−20 °C) for 3 minutes and in cold acetone (−20 °C) for 10 seconds. Samples were incubated with anti-Tub1 primary antibody (YOL1/34 Biorad), for 2 hours at room temperature and washed 3 times with 1% BSA -PBS. Samples were then incubated with secondary antibody (FITC-conjugated anti-rat antibody from Jackson ImmunoResearch Laboratories, preabsorbed) for 1 hour at room temperature in a dark chamber and washed 4 times with 1% BSA-PBS. DAPI (100 mg p-phenylenediamide + 10 ml PBS adjust to pH 8.0 ad DAPI 0.05ug/ml) was added in each sample and slides were closed. Images were acquired using DeltaVision Elite imaging system (Applied Precision) based on an inverted microscope (IX71; Olympus) with a camera (CoolSNAP HQ2; Photometrics) and a UPlanFL N100x oil immersion objective lens (NA 1.4, Olympus).

Spindle length was measured with a custom script (gift from Rosella Visintin’s lab) written in FIJI. This script computes the Laplacian of the maximum projection of fluorescence intensities, identifies the spindle using the RidgeDetection plugin on the cell selected by the user, and compute the spindle length. We measured budded cells, as identified by the DAPI signal. To include in the analysis only cells that had developed a mitotic spindle, we analyzed only those with a spindle longer than 1 μm. At least 100 cells per sample were included in the analysis.

### Flow Cytometry for cell cycle analysis

Cells were treated according to [6]. Briefly, exponentially growing cells fixed overnight in EtOH 70% were incubated in RNAse A solution (0.05M NaCitrate, 0.25 mg/ml RNAse A) for 3 hours at 37 °C then in proteinase K solution (0.05M TRIS-HCl pH8, 0.01M CaCl2, 0.25 mg/ml Proteinase K) overnight at 55 °C. After staining for 20 minutes in Sytox Green Solution (1μM Sytox Green, 0.05 M NaCitrate), samples were sonicated and acquired using attune cytofluorimeter (Thermo Scientific). Sytox fluorescence was acquired using BL1 laser. fcs files were analyzed using a customized R script using ggplot for visualization. Singlets were gated on SSC parameters. Cell cycle phases were discriminated based on Sytox Green signal (DNA content).

### Cell size measurements

For size analysis, ancestors were collected after 24 hours at 18 °C (i.e. first timepoint of evolutionary experiment-G0), while evolved strains at the end of the evolutionary experiment (Gf). Exponentially growing cells were fixed in EtOH 70% and stained with Sytox Green as described above. Samples were acquired at FACS (Macsquant) with the same voltage settings for all the samples. Doublet were excluded based on Sytox Green signal and size was approximated with FSC physical parameter.

### Sanger sequencing of selected genes

The indicated evolved strains were streaked on agar plates and incubated at 18 °C until colonies were visible (about 5 days). Then, colonies were picked and amplified on agar at 18 °C to make a glycerol stock. PCR to amplify genes to be sequenced was done directly from colonies or after defrosting and growing on agar at 18 °C: cells were boiled in 3 μl 20 mM NaOH for 10 minutes and directly added to PCR mix. The amplification products were then sequenced by Sanger sequencing.

### Live cells imaging experiment

To measure missegregation rate using single cell analysis, cells were grown overnight at 30 °C. The day after, cells were diluted at OD600 0.15, grown for 1 hour, supplied with α-factor [5 μg/ml], shifted at 18 °C for 1 hour, sonicated, and loaded on a microfluidic chamber (CellASIC), where they grew at 18 °C supplemented with α-factor for 30 minutes. Then, cells were released from G1 arrest, and they grew for 40 hours at 18 °C in the microfluidic chamber under the microscope. Missegregation was identified by following the positions of GFP-tagged chromosomes V after the first cell division or adaptation. Temperature was controlled as in [7] using a BOLD LINE Water-Jacket Top Stage Incubation System (OKOlab).

Time-lapse movies were recorded using a DeltaVision Elite imaging system (Applied Precision) based on an inverted microscope (IX71; Olympus), a UPlanFL N 60x (1.25 NA) oil immersion objective lens (Olympus) and a camera (Scientific CMOS Camera).

Reference images were taken every 15 minutes, while fluorescence images every hour. GFP was acquired using single bandpass filters (EX475/28 EM523/36) with 5 z-stack (0.85 μm), exposure time 0.105 s, and power lamp 10%. To evaluate any phototoxicity of the acquisition settings, cell cycle duration of excited and non-excited fields of *TUB2* strains was compared, and no difference was observed. Z-stacks were deconvolved with SoftWoRx software and then projected with maximum intensity projection with Fiji.

Cells were segmented and tracked using a custom version of phylocell [8], and missegregation was assessed semi-automatically using a MATLAB script based on the FindPeaks function.

### Whole-genome sequencing and analysis

#### DNA extraction

To extract DNA of ancestors or evolved populations (Gr and Gf), cells were inoculated in YPD at 30 °C or 18 °C, respectively, and grown for 1 or 4 days, respectively. Then, DNA was extracted from 10 ml of yeast culture at stationary phase. Cell wall was digested at 37 °C with 200 μl of SCE solution (1M Sorbitol, 0.1M NaCitrate, 0.06M EDTA, pH 7.0) supplemented with 2 mg/ml of zymolyase and 8 μl/ml of 2-mercaptoethanol for 30-60 minutes. Spheroplasts were lysed SDS solution (SDS 2%, 0.1M Tris/HCl pH 9.0, 0.05M EDTA) at 65 °C for 5 minutes. DNA were purified with standard NH4OAc/isopropanol precipitation and resuspended in 200 μl of water supplemented with 1 μl of RNAse 10 mg/ml. RNA was removed by overnight digestion at 37 °C. Following, DNA were purified with a second round of NH4OAc/Isopropanol precipitation and resuspended in water.

#### Sequencing and bam files creation

Sequencing was performed at Beijing Novogene Bioinformatics Technology Co., Ltd., using an Illumina high-throughput sequencer returning 2×150 bp paired-end reads.

FASTA files were trimmed using trimmomatic v0.36 [9], then aligned to reference sequence (sacCer3) using Burrows-Wheeler Aligner (BWA) v0.7.17 [10]. Then, duplicates were removed using samblaster v0.1.24 and sorted using samtools v1.9 [12]. Lastly, bam files were realigned around indels using Picard v2.19.0 [13] and GATK v3.8-1 [14]. Final bam files have an average depth of coverage of 50 reads. Notice that ancestors were clonal, while evolved populations were mixed.

#### Identification of single-nucleotide changes and short indels

We aimed at identifying single-nucleotide changes and short indels (less than ∼50 bp) filtering out variations of noisy genomic regions or specific of the genetic background (W303). To do so, we used IsoMut [15]. Briefly, this tool compares a set of isogenic sample sequences, and calls a variation when it is unique for one of the sequences. To identify variations in each of our target samples, we fed IsoMut with a set of 11 sequences: the target sample sequence, plus 10 technical replicates of wild-type yAC4179, which were used to remove W303-specific variations and noisy genomic regions. As output, IsoMut returns the variations on non-noisy regions that the sample does not share with the background set of 10 wild-type. The variations in all the samples (ancestors and evolved populations) are then joined in a unique database.

We then filtered the mutations, using a conservative approach. Using a custom written R script based on VariantAnnotation v1.28.11 [16], mutations were mapped to genomic regions. Following [17], we removed regions annotated in the SGD database as simple repeats, centromeric regions, telomeric regions, or LTRs (SGD project; ttp://downloads.yeastgenome.org/curation/chromosomal_feature/ SGD_features.tab, downloaded July 26, 2019). To identify mutations with good signal, we removed those identified by <5 reads, regardless of their allelic frequency, those with IsoMut cleanliness parameter <0.98, those with IsoMut score <0.21.

We analysed the mutations found in the two ancestors, to assess their isogenicity. They differ by 9 mutations (listed in Table S6), 5 of which are nonsynonymous.

To identify potentially adaptive mutations, we removed from the evolved strains the mutations inherited from their ancestor and those found in mitochondrial DNA. We then identified genes mutated in more than one population at the end of the experiment (Gf) by unique mutations resulting in non-synonymous amino acid changes. This definition gave a list of 10 genes (*PHO4, PHO81, KIP3, TUB1, TUB2, SPC98, ADE6, PRR2, YHR033W, YJL070C*). These genes form the core of the ‘recurrently mutated genes‘, genes whose mutations are potentially adaptive to cold or impaired microtubule polymerization. In the final list (genes shown in Table S1), we did not include *PRR2, YHR033W, ADE6* and *YJL070C* as they are present only twice in very low frequencies. Instead, we included *TUB4*, which was mutated twice with high frequency but with identical mutations, and for this reason originally not present in the list. Nevertheless, we decided to include it based on the fact that we had identified *SPC98*, another component of gamma-TuSC, which suggests mutations in gamma-TuSC being potentially adaptive.

Having defined the recurrently mutated genes, eventually, we kept track of all mutations that hit any of them in any population. 61 mutations (listed in Table S1) belong to this group. There are other 146 mutations which hit other genes and are not reported in Table S1: either they did not fulfill the conditions to be ‘recurrently mutated’ (unique mutations in more than one population), or were excluded (*YHR033W, YJL070C, PRR2, ADE6*) as explained above. Mutations of recurrently mutated genes have higher frequencies than the other mutations (median of mutations frequencies 0.218 and 0.086, respectively). In the manuscript, we focused our analysis on mutations hitting recurrently mutated genes.

### Analysis of aneuploidy

Depth of coverage was extracted from bam files using a custom script written in python that makes use of the “depth” function contained in the samtools package. Chromosomes were partitioned into bins of 10000 bp, and the bin-wise median was used for further analysis.

In order to avoid an influence of outliers (e.g. due to large peaks resulting from repeat elements), we removed values more than 20% below or above the median signal for each chromosome.

We noticed a pronounced bias of depth of coverage apparently related to chromosome length. Plotting chromosome median depth of coverage against chromosome length suggests that this relation can be approximated by an exponential of the form

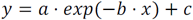

We fit each sample individually and divided the median depth for each bin by the value corresponding to the respective chromosome resulting from the fit in order to detrend the data. We used robust regression in order to avoid an influence on detrending due to the actual biological signal. Thus, clear aneuploidies are treated as outliers that do not distort the fit.

For individual chromosomes we noticed a further bias of depth of coverage towards the ends of chromosomes, resulting in a “smile” shape. In order to correct for this bias, we superposed the signal for all chromosomes, normalizing by median and length, and then fitting a quadratic function. The median of the corrected depth of coverage signal was used as a proxy for chromosomal copy number.

### Analysis of Structural Variations

In order to search for structural variations beyond the level of SNVs and small indels, we applied a set of different methods, as there is no single method that detects the full spectrum of possible variants with high sensitivity and specificity. The identification of variants is further complicated by the fact that mixed populations were sequenced.

We used approaches based on read pair (BreakDancer [18]), split read (Pindel [19]), and read count (CNVnator [20]). The read count method was strongly affected by the “smile” bias mentioned above. In order to remove the bias, we downsampled the bam files using samtools “view” proportional to the quadratic fit. To remove false positive calls and variants expected due to genetic differences between the strain used in the experiments (W303) and the reference genome (S288C), we filtered out variants shared by multiple wells using mutual overlap as a criterion for the identification of two variants. Promising candidates were checked by eye using IGV [21]. To convince ourselves of the sensitivity of our methods for discovering larger structural variants, we verified that the amplifications of *URA3* and *TRP1* were actually detected in the strains. However, all our candidates turned out to be either artifacts (in one case due to contamination by a human sequence) or already detected as indels using the basic alignment.

